# Enhanced mGluR1 function causes motor deficits and region-specific Purkinje cell dysfunction

**DOI:** 10.1101/2025.03.24.644877

**Authors:** Mohamed F. Ibrahim, Sevda Boyanova, Yin Chun Cheng, Clemence Ligneul, Rasneer S. Bains, Jason P. Lerch, Ed Mann, Peter L. Oliver, Esther B. E. Becker

**Affiliations:** Nuffield Department of Clinical Neurosciences, University of Oxford, Oxford, OX3 9DU, UK; Kavli Institute of Nanoscience Discovery, University of Oxford, Oxford, OX1 3QU, UK; Wellcome Centre for Integrative Neuroimaging, FMRIB, Nuffield Department of Clinical Neurosciences, University of Oxford, Oxford, OX3 9DU, UK; Mary Lyon Centre at MRC Harwell, Didcot, OX11 0RD, UK; Department of Medical Biophysics, University of Toronto, Toronto, ON, M5G 2C4, Canada; Department of Physiology, Anatomy and Genetics, University of Oxford, Oxford, OX1 3PT, UK; Mammalian Genetics Unit, MRC Harwell Institute, Didcot, OX11 0RD, UK

**Keywords:** Cerebellum, Grm1, mGluR1, Purkinje cell, ataxia, selective vulnerability

## Abstract

Spinocerebellar ataxias (SCAs) are autosomal dominantly inherited neurodegenerative disorders with no effective treatment. Aberrant signalling through the metabotropic glutamate receptor (mGluR1) has been implicated in several SCAs. However, whether disease is caused through decreased or increased mGluR1 signalling remains controversial. Here, we generate the first mouse model of enhanced mGluR1 function by introducing a gain-of-function mutation (p.Y792C) that causes SCA44 in the metabotropic glutamate receptor 1 *(Grm1*) gene. *Grm1* mutant mice recapitulate key pathophysiological aspects of SCA, including progressive motor deficits, altered climbing fibre innervation and perturbed Purkinje cell spontaneous activity. We report that changes in synaptic innervation and intrinsic Purkinje cell activity upon overactive mGluR1 signalling manifest in a lobule- and disease-stage-specific manner. Our findings demonstrate that enhanced mGluR1 function is a direct and specific driver of Purkinje cell dysfunction and pathology and provide a mechanism for understanding the selective vulnerability of different Purkinje cell populations in SCA.

## Introduction

The spinocerebellar ataxias (SCAs) are a genetically and clinically complex group of autosomal dominant neurodegenerative diseases that primarily affect the cerebellum and its associated pathways^1,2^. SCAs are characterised by a progressive loss of balance and coordination accompanied by slurred speech, difficulties in swallowing, and oculomotor abnormalities. SCAs are heterogeneous, and to date more than 40 genetically-defined subtypes have been identified. The overall rarity of SCAs, together with the diverse genes and implicated pathways that cause the disease, renders the feasibility of finding therapeutics for individual subtypes challenging^3^. Therefore, a deeper understanding of the disease mechanisms that might be shared by distinct SCA subtypes is critical to advancing treatment options.

Emerging evidence indicates that aberrant signalling through the metabotropic glutamate receptor type 1 (mGluR1) is a central mechanism in many SCAs^4,5^. mGluR1 is highly expressed in Purkinje cells and plays a crucial role in the development, synaptic wiring, spontaneous activity and plasticity of Purkinje cells^5–8^. Intriguingly, both reduced and increased mGluR1 function have been implicated in cerebellar ataxia in human patients and genetically engineered mouse models. Autoantibodies against mGluR1 and downstream signalling molecules have been identified in autoimmune ataxias^9^, and homozygous loss-of-function mutations in *the GRM1* gene encoding mGluR1 are associated with autosomal recessive spinocerebellar ataxia 13 (SCAR13)^10,11^. Downregulation of mGluR1 and associated signalling molecules has been reported in some mouse models of SCAs including SCA1, SCA2 and SCA3^12–16^. However, it remains controversial whether these changes are an early, pathogenic mechanism in SCA or whether they represent a later, potentially compensatory consequence to maintain calcium homeostasis. In support of the latter, prolonged and thus enhanced mGluR1 currents were observed in the conditional SCA1 [82Q] mouse model, which has a more moderate phenotype and likely represents an early disease stage of SCA1^17^. mGluR1 signalling was also found to be greatly amplified early in disease in a mouse model of SCA2^18^. In support of the hypothesis that overactive mGluR1 signalling causes cerebellar ataxia, gain-of-function mutations in the TRPC3 cation channel that is activated by mGluR1 cause cerebellar ataxia in the *Moonwalker* (*Mwk*) mouse mutant and patients with SCA41^19,20^. Similarly, in a mouse model of SCA14 mutant PKCy fails to inhibit TRPC3 and, as a consequence, mGluR1-mediated currents are increased^21^. Based on these and other studies, it has been postulated that a positive feedback loop between enhanced mGluR1 signalling and elevated calcium levels in Purkinje cells might be a common driver of Purkinje cell dysfunction and loss in SCA^18^. However, so far evidence has been lacking on whether enhanced mGluR1 can directly cause Purkinje cell dysfunction and disease.

In this study, we present a mouse model of enhanced mGluR1 function that allowed us to investigate for the first time the direct impact of altered mGluR1 activity on Purkinje cell and cerebellar function. We previously reported the first dominant mutations in the *GRM1* gene causing SCA44 in two families with adult-onset cerebellar ataxia and degeneration^22^. Using an mGluR1 reporter assay in heterologous cell lines, we showed that the SCA44 patient *GRM1* mutations (p.Y262C and p.Y792C) behaved as gain-of-function *in vitro*^22^. This is consistent with a recent pharmacological study that demonstrated enhanced constitutive activity of the SCA44 p.Y792C mutation in calcium mobilisation studies in HEK293 cells^23^. While these findings support a role for enhanced mGluR1 activity in SCA44, the effect of the gain-of-function *GRM1* mutations on the cerebellum remained unexplored, and thus the pathophysiological mechanisms underlying SCA44 are not well understood. We therefore developed a *Grm1* mutant mouse model harbouring the SCA44 p.Y792C mutation and demonstrate that this mutation results in enhanced mGluR1 signalling in Purkinje cells. Behaviourally, heterozygous *Grm1*^Y792C/+^ mutant mice recapitulate the late-onset ataxic phenotype observed in human SCA44 patients, whereas homozygous *Grm1*^Y792C/Y792C^ mutant mice develop progressive cerebellar ataxia from three months of age. Moreover, *Grm1* mutants display altered climbing fibre-Purkinje cell synaptic connectivity and aberrant Purkinje cell activity, which are both key pathological hallmarks of SCAs. Notably, we also show that the observed structural and functional deficits are region- and disease-stage-dependent. Together, our study demonstrates that enhanced mGluR1 function is a direct driver of selective Purkinje cell dysfunction leading to cerebellar ataxia.

## Results

### Introduction of a SCA44 patient mutation into mouse *Grm1* results in enhanced mGluR1 activity in Purkinje cells

To investigate the pathophysiological mechanisms underlying SCA44 and, more broadly, to understand the role of enhanced mGluR1 signalling in cerebellar function and disease, we generated a mouse model harbouring the mGluR1 p.Y792C SCA44 patient gain-of-function mutation^22^. Tyr792 is located within the sixth helix transmembrane domain of mGluR1 (Fig. 1A), a critical region for receptor activation^24^. To generate the SCA44 patient mutation in the mouse, we employed CRISPR-Cas9-mediated gene editing and introduced an A-to-G base pair substitution in the endogenous mouse *Grm1* gene, resulting in the mGluR1 p.Y792C missense mutation. The successful generation of both heterozygous (*Grm1*^Y792C/+^) and homozygous (*Grm1*^Y792C/Y792C^) mutant animals was validated using Sanger sequencing (Fig. 1B).

**Figure 1.**
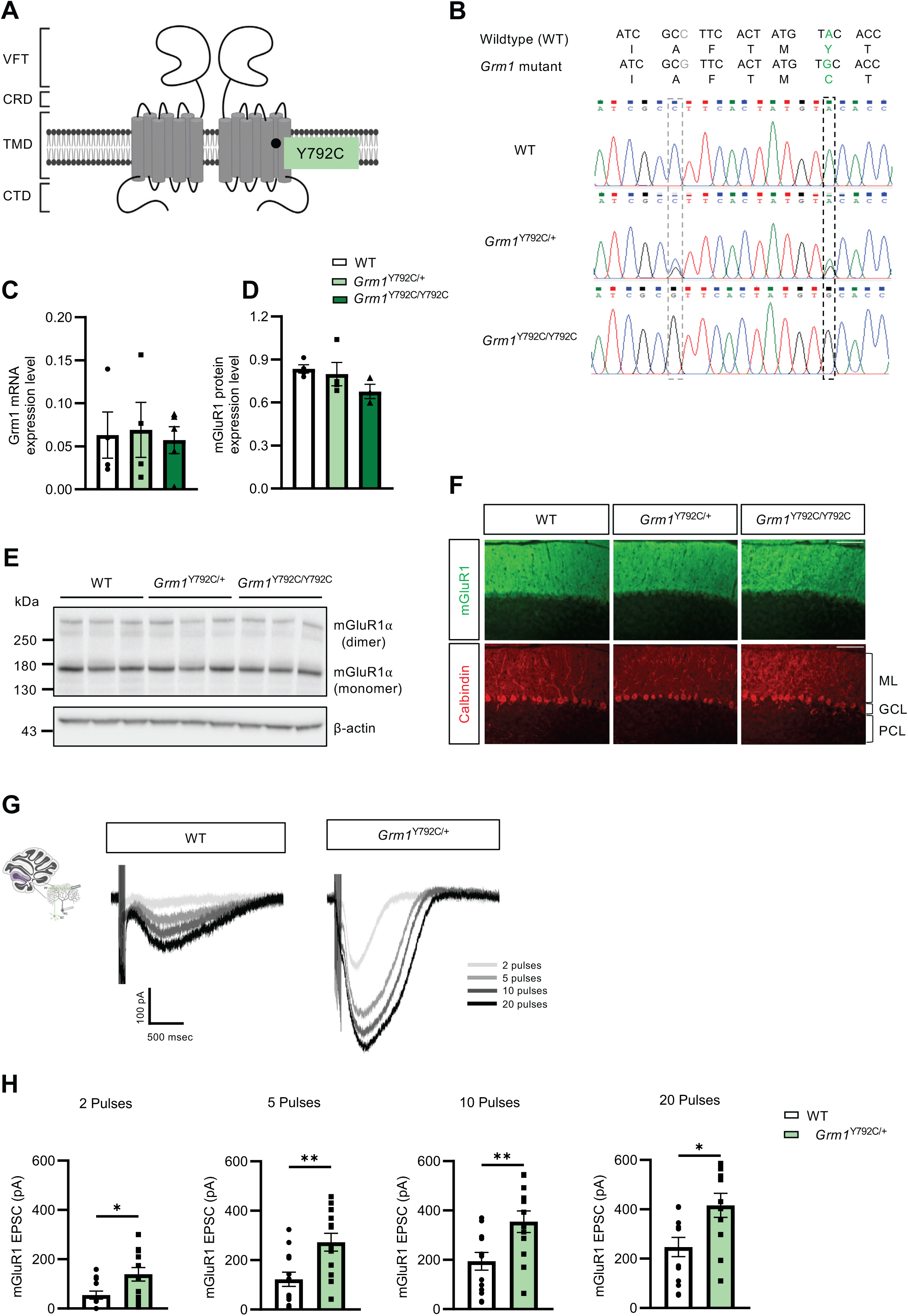
Characterisation of mGluR1 expression and synaptic transmission in *Grm1* mutant cerebellum. **(A)** Schematic representation of the SCA44 p.Y792C mutation within the mGluR1 receptor. VFT: venus fly trap domain; CRD: cysteine-rich domain; TMD: transmembrane domain (TMD); CTD: intracellular C-terminal domain. **(B)** Sanger sequencing confirms successful Cas9-mediated base-pair editing. In addition to the SCA44 point mutation (A>C; p.Tyr792Cys), a silent base pair substitution (C>G; p.Ala782Ala) was created in the nearby PAM site to prevent re-cutting by the Cas9 enzyme. **(C)** Quantification of relative *Grm1* mRNA expression levels in three-month-old wildtype (WT) and *Grm1* mutant mouse cerebellum by RT-qPCR. Expression calculated as 2^-ΔCT^, relative to housekeeper gene *Gapdh.* Three technical replicates were included in all experiments. Datapoints represent individual animals. WT vs. *Grm1*^Y792C/+^: *P*>0.9999, WT vs. *Grm1*^Y792C/Y792C^: *P*>0.9999, *Grm1*^Y792C/+^ vs *Grm1*^Y792C/Y792C^*: P*>0.9999. n=4-5 animals per genotype. One-way ANOVA followed by Bonferroni’s multiple comparison test. Error bars represent SEM. **(D)** Quantification of relative mGluR1 protein expression levels in three-month-old WT and *Grm1* mutant cerebellum. Protein levels are normalised to β-actin. Datapoints represent individual animals. WT vs *Grm1*^Y792C/+^: *P*>0.9999, WT vs *Grm1*^Y792C/Y792C^: *P*=0.3237, *Grm1*^Y792C/+^ vs *Grm1*^Y792C/Y792C^: *P*=0.6007. n=3-4 mice per group. One-way ANOVA followed by Bonferroni’s multiple comparisons. Error bars represent SEM. **(E)** Representative immunoblotting images showing expression of mGluR1 in the cerebellum of three-month-old WT and *Grm1* mutant littermates (n=3 mice per group). β-actin was used as loading control. Uncropped images with molecular weight markers are available in Supplementary Figure 6. **(F)** Representative immunostaining against mGluR1 (red) and calbindin (green) in three-month-old WT and *Grm1* mutant cerebellum. ML: molecular layer; PCL: Purkinje cell layer; GCL: granule cell layer. Scale bar: 100 μm. **(G)** Representative traces of parallel fibre-evoked slow mGluR1 EPSCs in WT and *Grm1*^Y792C/+^ Purkinje cells in lobule III of 5-7-week-old mice measured in extracellular solution containing antagonists for AMPA/kainite receptors, GABA(A) receptors and glutamate transporters. **(H)** Quantification of average-peak slow mGluR1 EPSC for increasing pulses (2, 5, 10 and 20 pulses) of 200-Hz stimulations. n=11-14 cells per genotype. WT vs *Grm1*^Y792C/+^: 2 pulses: *P*=0.0133, 5 pulses: *P*=0.0034; 10 pulses: *P*=0.0091; 20 pulses: *P*=0.0134. Statistical significance was determined by Mann-Whitney or unpaired t-test. \**P*<0.05, \*\**P*<0.01. Error bars represent SEM. For full statistical data, see Supplementary Table 2.

We first assessed whether the introduced missense mutation would affect mGluR1 expression in the cerebellum by quantifying mutant *Grm1* mRNA and mGluR1 protein levels. We did not observe any significant changes in *Grm1* mRNA expression (Fig. 1C) or mGluR1 protein levels in the cerebellum (Fig 1D, E) in either heterozygous or homozygous *Grm1* mutant mice compared to wildtype (WT) littermates. The mGluR1 receptor is highly expressed in Purkinje cell dendrites^25^, and we found that the dendritic localisation of mGluR1 in Purkinje cells was not altered upon the introduction of the Y792C mutation, as shown by immunostaining (Fig. 1F).

Next, we examined the functional impact of the Y792C mutation on mGluR1-mediated signalling. mGluR1 activation is responsible for the slow excitatory postsynaptic current (EPSC) at the parallel fibre (PF)-Purkinje cell synapse^26^. We recorded slow EPSCs in adult Purkinje cells from WT and *Grm1*^Y792C/+^mutants in response to an increased number of pulses delivered in 200-Hz trains to the innervating PFs (Fig. 1G). We found that mutant *Grm1*^Y792C/+^ Purkinje cells exhibited significantly larger slow EPSCs than WT Purkinje cells (*P*=0.0133 (2 pulses); *P*=0.0034 (5 pulses), *P*=0.0091 (10 pulses); *P*=0.0124 (20 pulses)) (Fig. 1G, H). Together, these results demonstrate that the p.Y792C mutation does not affect the localization of mGluR1 but causes enhanced mGluR1 function in Purkinje cells, consistent with a dominant gain-of-function mechanism.

### *Grm1* mutant mice display a progressive decline in motor coordination

To investigate whether the introduced SCA44 mutation and resulting enhanced mGluR1 function affects motor function, we carried out a longitudinal behavioural study in *Grm1* mutant animals compared to WT littermates from three to 18 months of age, representing young adulthood until old age. We assessed both heterozygous and homozygous *Grm1* mutants, as we predicted that the latter could serve as an accelerated disease model^27^.

First, we assessed the gait of the mutant mice using the MouseWalker system that allows the high-resolution tracking of locomotion features such as limb positioning during walking^28^. Starting at six months of age, *Grm1*^Y792C/Y792C^ mutant mice had a significantly larger front limb stride width compared to *Grm1*^Y792C/+^ mice (*P*=0.0008), which became worse with increasing age compared to WT and *Grm1*^Y792C/+^ mice (Fig. 2A). No statistical difference was detected at 18 months of age, likely reflecting a reduction in the experimental cohort size due to aging. When we examined motor impairments during walking across a horizontal ladder using the Locotronic apparatus, *Grm1*^Y792C/Y792C^ mice displayed increased front limb errors from three months of age (*P*<0.0001), which worsened progressively with increasing age of the tested animals (Fig. 2B). *Grm1*^Y792C/+^ animals did not show any impairments during the testing period.

**Figure 2.**
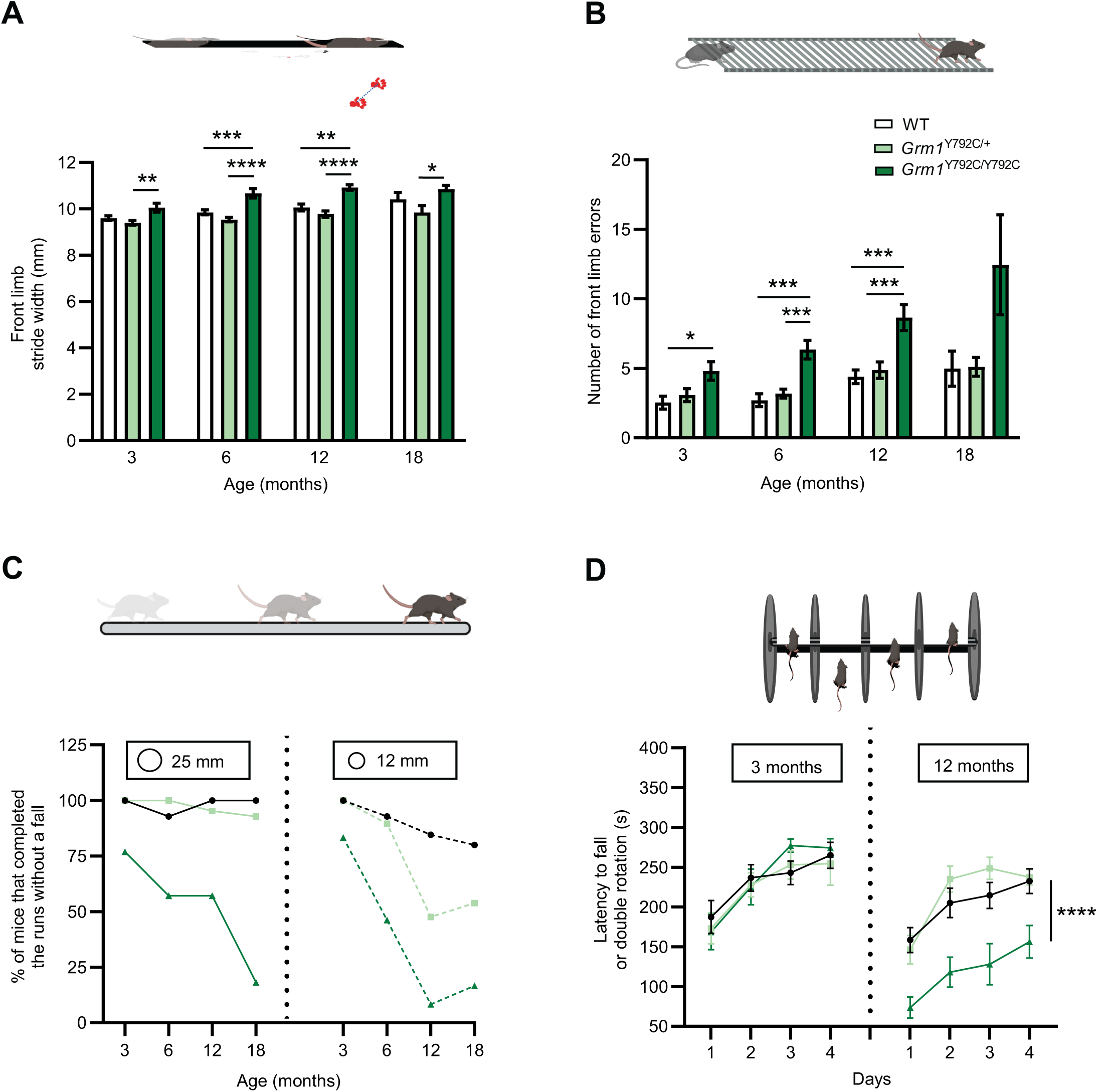
*Grm1* mutant mice develop progressive loss of motor performance and coordination. **(A)** Quantification of the front limb stride width, *i.e*., the distance between the right and left front limb placement, assessed using the MouseWalker system. 3 months: WT vs *Grm1*^Y792C/+^: *P*=0.4787, WT vs *Grm1*^Y792C/Y792C^: *P*=0.0707, *Grm1*^Y792C/+^ vs *Grm1*^Y792C/Y792C^: *P*=0.0020; 6 months: WT vs *Grm1*^Y792C/+^: *P*=0.2273, WT vs *Grm1*^Y792C/Y792C^: *P*=0.0008, *Grm1*^Y792C/+^ vs *Grm1*^Y792C/Y792C^: *P*<0.0001; 12 months: WT vs *Grm1*^Y792C/+^: *P*=0.3509, WT vs *Grm1*^Y792C/Y792C^: *P*=0.0012, *Grm1*^Y792C/+^ vs *Grm1*^Y792C/Y792C^: *P*<0.0001; 18 month: WT vs *Grm1*^Y792C/+^: *P*=0.4292, WT vs *Grm1*^Y792C/Y792C^: *P*=0.6214, *Grm1*^Y792C/+^ vs *Grm1*^Y792C/Y792C^: *P*=0.0201. Statistical significance was determined by one-way ANOVA followed by Tukey’s multiple comparison test. For n-numbers, see Supplementary Table 1. **(B)** Quantification of the number of front limb errors using the Locotronic apparatus. 3 months: WT vs *Grm1*^Y792C/+^: *P*>0.9999, WT vs *Grm1*^Y792C/Y792C^: *P*=0.0468, *Grm1*^Y792C/+^ vs *Grm1*^Y792C/Y792C^: *P*=0.0992; 6 months: WT vs *Grm1*^Y792C/+^: *P*>0.9999, WT vs *Grm1*^Y792C/Y792C^; *P*=0.0002, *Grm1*^Y792C/+^ vs *Grm1*^Y792C/Y792C^: *P*=0.0007; 12 months: WT vs *Grm1*^Y792C/+^: *P*=0.8573 , WT vs *Grm1*^Y792C/Y792C^: *P*=0.0003, *Grm1*^Y792C/+^ vs *Grm1*^Y792C/Y792C^: *P*=0.0008; 18 months: WT vs *Grm1*^Y792C/+^: P>0.9999, WT vs *Grm1*^Y792C/Y792C^: *P*=0.0093, *Grm1*^Y792C/+^ vs *Grm1*^Y792C/Y792C^: *P*=0.0526. Statistical significance was determined by one-way ANOVA followed by Tukey’s multiple comparison test or Kruskal-Wallis test followed by Dunn’s multiple comparison test. For n-numbers, see Supplementary Table 1. **(C)** Percentage of WT and *Grm1* mutant mice that have completed runs on the balance beam without a fall assessed on a wide (25 mm) and narrow (12.5 mm) beam. For n-numbers, see Supplementary Table 1. **(D)** Motor coordination represented as latency to fall or double rotation on an accelerating Rotarod of naïve female WT and *Grm1* mutant mice assessed at 3 and 12 months of age. 3 months: Genotype: F_(2,27)_=0.1120, *P*=0.8495; Day: F_(2.007,54.20)_=25.34, *P*<0.0001; 12 months: Genotype: F_(2,30)_=15.66, *P*<0.0001; Day: F_(2.642,79.27)_=25.38, *P*<0.0001. Statistical significance was determined by two-way ANOVA followed by Tukey’s *post hoc* test. Error bars represent SEM. \**P*<0.05, \*\**P*<0.01, \*\*\**P*<0.001, \*\*\*\**P*<0.0001. For n-numbers, see Supplementary Table 1. For full statistical data, see Supplementary Table 2.

We also assessed motor coordination and balance of the animals using a horizontal balance beam walking task with two beams of varying diameter. *Grm1*^Y792C/+^ mice performed to a similar level to WT mice on the wider beam (25 mm) at all ages of testing, with most animals able to traverse the beam without a fall. In contrast, the percentage of *Grm1*^Y792C/Y792C^ mice that successfully crossed the beam without a fall decreased markedly with increasing age (Fig. 2C). On the narrower beam (12.5 mm), *Grm1*^Y792C/Y792C^ mice showed impairments as early as three months of age, while deficits in *Grm1*^Y792C/+^ mice were evident from 12 months of age.

To investigate if there was any sex difference in the impact of motor coordination, we also examined gait, limb errors, and balance in a cohort of naïve 12-month-old female mutant mice. Similar to males, female *Grm1*^Y792C/Y792C^ mice exhibited a larger front limb stride width compared to WT (*P*=0.0029) (Supplementary Fig. 1A). Moreover, none of the female *Grm1*^Y792C/Y792C^ mice were able to traverse the wider balance beam without a fall (Supplementary Fig. 1B). These results show that the Y792C mGluR1 mutation affects motor coordination in both sexes.

Next, we examined aspects of motor coordination and motor learning in *Grm1* mutant mice using an accelerating Rotarod. No differences in the time taken to fall from the Rotarod, a measure of motor performance, were observed between naïve *Grm1* mutant mice and their WT littermates at three months of age (F_(2,27)_=0.1120; *P*=0.8495) (Fig. 2D). However, at 12 months of age, *Grm1*^Y792C/Y792C^ mice showed a significantly decreased motor performance compared to *Grm1*^Y792C/+^ and WT animals across all four days of testing (F_(2,30)_=15.66; *P*<0.0001) (Fig. 2D). In contrast, motor learning across consecutive testing days was not affected in the mutant animals, and they showed significant improvements over the testing days (day 1 vs day 4; WT: *P*=0.0045; *Grm1*^Y792C/+^, *P*=0.0017; *Grm1*^Y792C/Y792C^: *P*=0.0436) (Fig. 2D).

Taken together, our results show that mGluR1 gain-of-function results in a dose-dependent and progressive decline of motor coordination and balance in *Grm1* mutant mice, with homozygous *Grm1*^Y792C/Y792C^ mice showing deficits from an early age, and *Grm1*^Y792C/+^ mutants displaying a mild and later-onset phenotype. These findings are consistent with the progressive, late-onset ataxia described in human SCA44 patients that carry heterozygous *GRM1* mutations^22^.

### *Grm1* mutant mice are not impaired in non-cerebellar behaviours

In addition to its role in motor control, mGluR1 signalling has been implicated in other important brain functions including regulating spatial working memory^29,30^, anxiety-like behaviour^31,32^, sensorimotor gating^33^, and fear conditioning^34,35^. We therefore investigated these specific behaviours in both heterozygous and homozygous *Grm1* mutant mice.

First, we assessed short-term spatial working memory in WT and *Grm1* mutant mice using a Y-maze alternation test. In the initial habituation phase of the trial when only two arms were available, the total distance travelled was similar between all genotypes, indicating similar levels of general activity (Supplementary Fig. 2A). In the test phase, when all arms of the Y-maze were open for exploration, mice of all genotypes spent a similar proportion of their time in the novel arm compared to the total time spent in the novel and familiar arms; this finding was consistent at all four timepoints tested, suggesting that short-term spatial working memory is unaltered in both *Grm1* mutants (Supplementary Fig. 2B).

Next, to evaluate *Grm1* mutant mice for anxiety-like behaviour, we applied three independent tests that compare time spent in open versus closed areas of the apparatus. As before, the total distance travelled in these tests was similar between all three genotypes, suggesting that the results from these tests were not influenced by differences in general activity (Supplementary Fig. 2C, E, G). In the open field and the light/dark box, the percentage of time spent in the open central zone and in the dark zone, an indication of non-anxious behaviour, was similar between the WT and *Grm1* mutant mice (Supplementary Fig. 2D, F). Some differences were observed between WT and *Grm1*^Y792C/Y792C^ animals in the zero maze (Supplementary Fig. 2H), but these were inconsistent, with the mutants only at three and 12 months spending more time in the open arm and less time in the closed arm than WT. Overall, these results suggest that anxiety-like behaviour is not affected in *Grm1* mutant mice.

In the fear conditioning test, where freezing behaviour is measured as an index of context-specific fear memory, *Grm1* mutant mice exhibited similar freezing levels to WT mice before and after the conditioning tone (Supplementary Fig. 3A). We also assessed sensorimotor gating by measuring the prepulse inhibition (PPI) of an auditory startle response. PPI levels increased with increasing prepulse intensity as expected, although there were no significant differences between the three genotypes (Supplementary Fig. 3B). Together, our behavioural results demonstrate that the gain-of-function Y792C mutation in mGluR1 causes behavioural deficits that are linked specifically to motor-and balance-related cerebellar dysfunction, and that enhanced mGluR signalling does not significantly affect the other behaviours tested here.

### Structural changes in the *Grm1* mutant cerebellum

Given the observed cerebellar behavioural deficits in *Grm1* mutant mice, we next investigated whether mutant animals display any structural changes in the cerebellum. We found that the gross cerebellar morphology of mutant mice was indistinguishable from WT mice, even at late age (Supplementary Fig. 4). To quantitatively assess potential degenerative changes, we performed immunostaining for calbindin, a marker for Purkinje cells, at both early symptomatic (three month) and late progressive (15 month) disease stage. We quantified the number of Purkinje cells and the thickness of the molecular layer in WT and *Grm1* mutant cerebellum. Measurements were taken across different lobules of the cerebellum (III, V and X), as anterior and posterior cerebellar regions are known to be differentially affected in cerebellar disease models^36,37^. We found no significant changes in the thickness of the molecular layer or the number of Purkinje cells in the *Grm1* mutant (both *Grm1*^Y792C/+^ and *Grm1*^Y792C/Y792C^) cerebellum (Fig. 3B-E). However, a subtle but significant reduction of the Purkinje cell soma area was found in the *Grm1* mutant cerebellum in the anterior lobules (III and V) at three months of age (Fig. 3F). By 15 months, reduced Purkinje cell soma size was evident across all measured lobules and in both *Grm1*^Y792C/+^ mice and *Grm1*^Y792C/Y792C^ mice (Fig. 3G). These results suggest that enhanced mGluR1 function does not alter gross cerebellar architecture or cause Purkinje cell loss during the observed time course. However, enhanced mGluR1 function causes subtle but progressive Purkinje cell soma atrophy, with cells in the anterior cerebellum being specifically vulnerable in both heterozygous and homozygous mutants.

**Figure 3.**
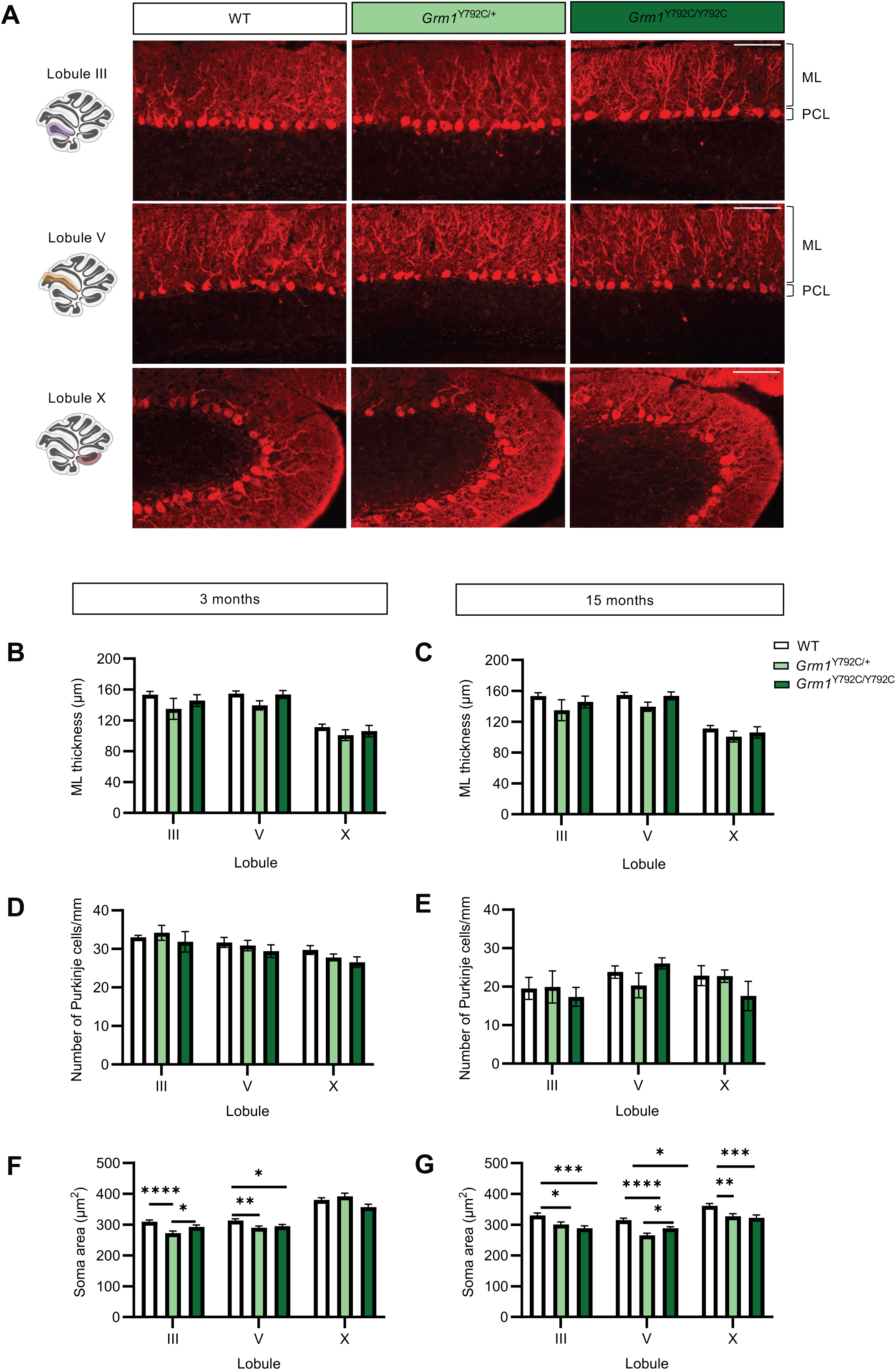
Adult *Grm1* mice exhibit lobule-specific Purkinje cell soma atrophy without cell loss or dendritic atrophy. **(A)** Representative immunostaining for the Purkinje cell marker calbindin in lobules III, V and X of 15-month-old WT and *Grm1* mutant cerebellum. ML: molecular layer; PCL: Purkinje cell layer. Scale bar: 100 μm. **(B, C)** Quantification of the molecular layer (ML) thickness in different lobules (III, V and X) in three- and 15-month-old WT and *Grm1* mutant cerebellum. No statistical differences are found. **(D, E)** Quantification of Purkinje cell density in different lobules (III, V and X) in three- and 15-month-old WT and *Grm1* mutant cerebellum. No statistical differences are found. **(F)** Quantification of Purkinje cell soma area in different lobules (III, V and X) in three-month-old WT and *Grm1* mutant cerebellum. Lobule III: WT vs *Grm1*^Y792C/+^: *P*<0.0001, WT vs *Grm1*^Y792C/Y792C^; *P*=0.107, *Grm1*^Y792C/+^ vs *Grm1*^Y792C/Y792C^: *P*=0.0504; lobule V: WT vs *Grm1*^Y792C/+^: *P*=0.0093, WT vs *Grm1*^Y792C/Y792C^; *P*=0.0501, *Grm1*^Y792C/+^ vs *Grm1*^Y792C/Y792C^: *P*>0.9999; lobule X: WT vs *Grm1*^Y792C/+^: *P*>0.9999, WT vs *Grm1*^Y792C/Y792C^; *P*=0.1147, *Grm1*^Y792C/+^ vs *Grm1*^Y792C/Y792C^: *P*=0.1165. **(G)** Quantification of Purkinje cell soma area in different lobules (III, V and X) in 15-month-old WT and *Grm1* mutant cerebellum. Lobule III: WT vs *Grm1*^Y792C/+^: *P*=0.0364, WT vs *Grm1*^Y792C/Y792C^; *P*=0.0009, *Grm1*^Y792C/+^ vs *Grm1*^Y792C/Y792C^: *P*=0.8773; lobule V: WT vs *Grm1*^Y792C/+^: *P*<0.0001, WT vs *Grm1*^Y792C/Y792C^; *P*=0.0227, *Grm1*^Y792C/+^ vs *Grm1*^Y792C/Y792C^: *P*=0.037; lobule X: WT vs *Grm1*^Y792C/+^: *P*=0.0137, WT vs *Grm1*^Y792C/Y792C^; *P*=0.0003, *Grm1*^Y792C/+^ vs *Grm1*^Y792C/Y792C^: *P*>0.9999. n=3-4 animals for each genotype. Statistical significance was determined by one-way ANOVA followed by Bonferroni’s multiple comparison test or Kruskal-Wallis test followed by Dunn’s multiple comparison test. Error bars represent SEM. \**P*<0.05, \*\**P*<0.01, \*\*\**P*<0.001, \*\*\*\**P*<0.0001. For full statistical data, see Supplementary Table 2.

To investigate changes to the cerebellar structure in greater detail, we next employed MRI imaging. We chose to image the *Grm1*^Y792C/Y792C^ animals compared to WT littermates at two different timepoints, as only the homozygous mutants were showing clear progressive ataxic phenotypes from an early age (Fig. 2). At an early symptomatic stage (2.5 months), we observed regional atrophy in the *Grm1* mutant cerebellum, affecting both the cerebellar cortex and the deep cerebellar nuclei (Fig. 4A). In the cerebellar cortex, the affected regions were confined to the lateral hemispheres with significant volume reductions in Crus1 and simplex lobule gray and white matter. Consistent with our histological findings, the atrophy was more pronounced in the anterior cerebellum. The interposed nucleus was also significantly smaller in the mutant mice compared to WT littermates. To evaluate whether the cerebellar atrophy worsened with disease progression, we imaged the mice again at six months of age. While the same cerebellar regions remained atrophied, we did not find an increase in the severity of atrophy or additional regions affected. However, we cannot rule out that cerebellar atrophy continues over a longer period of time. Together, our findings show that enhanced mGluR1 function results in cerebellar atrophy with anterior regions of the cerebellum affected the most.

**Figure 4.**
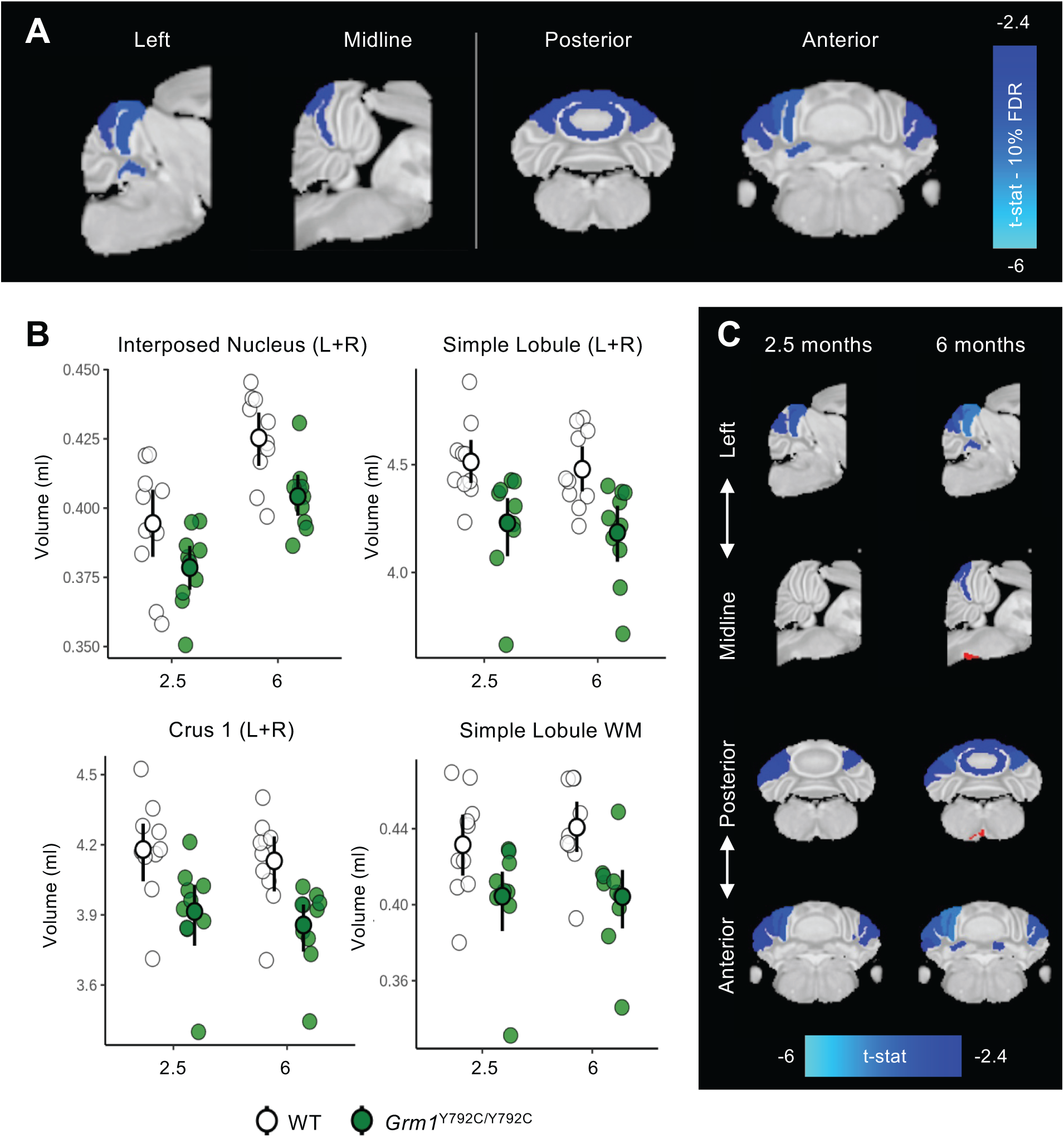
MRI imaging reveals regional atrophy in the *Grm1* mutant cerebellum. **(A)** *Grm1*^Y792C/Y792C^ animals present regional atrophy in the cerebellum. **(B)** Mean bilateral volumetric changes between WT (white) and mutant (green) are detected at 2.5 and 6 months of age for the interposed nucleus, Crus 1, the simple lobule and its associated white matter. **(C)** Volumetric changes in the cerebellum observed at 6 months already exist at 2.5 months.

### Region- and disease stage-specific climbing fibre deficits in *Grm1* mutant cerebellum

mGluR1 signalling is implicated in the spatial organization of parallel fibre (PF) and climbing fibre (CF) synapses onto the Purkinje cells^7^. Moreover. altered CF extension along the Purkinje cell dendritic tree (CF territory) has been observed in several ataxic mouse models including those linked to aberrant mGluR1 signalling^38,39^. To investigate whether increased mGluR1 activity can directly cause changes in PF and CF innervation, we performed immunostaining for vesicular glutamate transporter 1 (VGluT1), a marker for PF terminals, and VGluT2, a marker for CF terminals in *Grm1* mutant animals at early (three months) and late stage (15 months) of disease. We found that at three months of age, the extend of CF territory was significantly reduced in the anterior lobule III in *Grm1*^Y792C/Y792C^ mice compared to WT littermates (*P*=0.0012) (Fig. 5A, C). In contrast, CF territory in the posterior lobule X was similar in WT and *Grm1* mutant cerebellum (Fig. 5B, C). At 15 months of age, CF territory in the *Grm1*^Y792C/Y792C^ cerebellum was significantly reduced in both lobule III (*P*=0.0025) and lobule X (*P*=0.0239) (Fig. 5A-D). No significant reduction in CF territory in was observed in either lobule III or X in the *Grm1*^Y792C/+^ cerebellum, even at late stage of disease.

**Figure 5.**
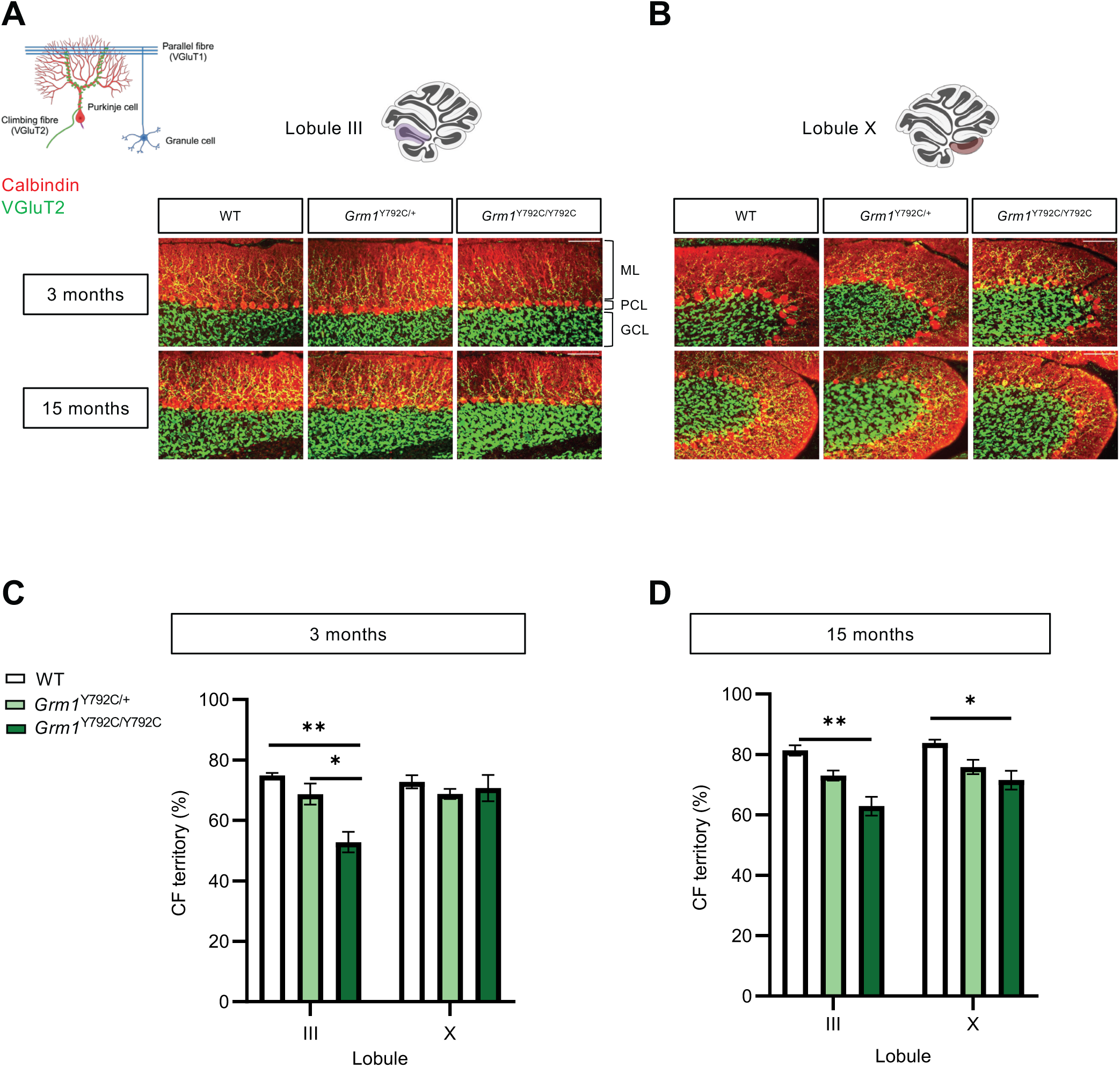
Lobule- and age-specific reduction in climbing fibre synapse innervation in *Grm1* mutant cerebellum. **(A, B)** Representative immunostaining for the climbing fibre synapse marker VGluT2 (green) and the Purkinje cell marker calbindin (red) in the anterior lobule III and posterior lobule X from three-month-old (top) and 15-month-old (bottom) WT and *Grm1* mutant cerebellum. ML: molecular layer; PCL: Purkinje cell layer; GCL: granule cell layer. Scale bar: 100 μm. **(C)** Quantification of the percentage of the climbing fibre (CF) territory in lobules III and X in three-month-old WT and *Grm1* mutant cerebellum. Lobule III: WT vs *Grm1*^Y792C/+^: *P*=0.4057, WT vs *Grm1*^Y792C/Y792C^; *P*=0.0018, *Grm1*^Y792C/+^ vs *Grm1*^Y792C/Y792C^: *P*=0.013; lobule X: WT vs *Grm1*^Y792C/+^: *P*=0.8761, WT vs *Grm1*^Y792C/Y792C^; *P*>0.9999, *Grm1*^Y792C/+^ vs *Grm1*^Y792C/Y792C^: *P*>0.9999. n=3-4 animals for each genotype. Statistical significance was determined by one-way ANOVA followed by Bonferroni’s multiple comparison test. Error bars represent SEM. \*\**P*<0.01. **(D)** Quantification of the percentage of the climbing fibre (CF) territory in lobules III and X in 15-month-old WT and *Grm1* mutant cerebellum. Lobule III: WT vs *Grm1*^Y792C/+^: *P*=0.1920, WT vs *Grm1*^Y792C/Y792C^; *P*=0.0037, *Grm1*^Y792C/+^ vs *Grm1*^Y792C/Y792C^: *P*=0.0729; lobule X: WT vs *Grm1*^Y792C/+^: *P*=0.2424, WT vs *Grm1*^Y792C/Y792C^; *P*=0.0358, *Grm1*^Y792C/+^ vs *Grm1*^Y792C/Y792C^: *P*=0.8204. n=3-4 animals for each genotype. Statistical significance was determined by one-way ANOVA followed by Bonferroni’s multiple comparison test. Error bars represent SEM. \**P*<0.05, \*\**P*<0.01. For full statistical data, see Supplementary Table 2.

We did not observe any prominent changes in the VGluT1 immunoreactivity or localisation in *Grm1* mutant cerebellum (Supplementary Fig. 5), suggesting that PF-Purkinje cell synapses remain unaffected by mGluR1 gain-of-function. Together, these results indicate that enhanced mGluR1 signalling causes a progressive reduction in CF innervation that starts in the anterior cerebellum before extending to posterior regions.

### Enhanced mGluR1 activity causes early disruption of Purkinje cell spontaneous activity

Purkinje cells fire spontaneous action potentials at high frequencies, and this spontaneous activity is commonly found to be altered in cerebellar diseases^40^. To investigate whether enhanced mGluR1 activity affects the spontaneous activity of Purkinje cells, we performed cell-attached recordings in acute cerebellar slices from both heterozygous and homozygous *Grm1* mutant mice and WT littermates. We recorded spontaneous activity both at an early disease stage (2.5 months) and at a late timepoint (15-16 months), when the disease has progressed in *Grm1*^Y792C/Y792C^ mice and subtle motor deficits are present in *Grm1*^Y792C/+^ animals (Fig. 2C). We observed three patterns of Purkinje cell spontaneous activity: continuous regular tonic firing, bursting or intermittent irregular firing, and silent cells with no firing activity (Fig. 6A).

**Figure 6.**
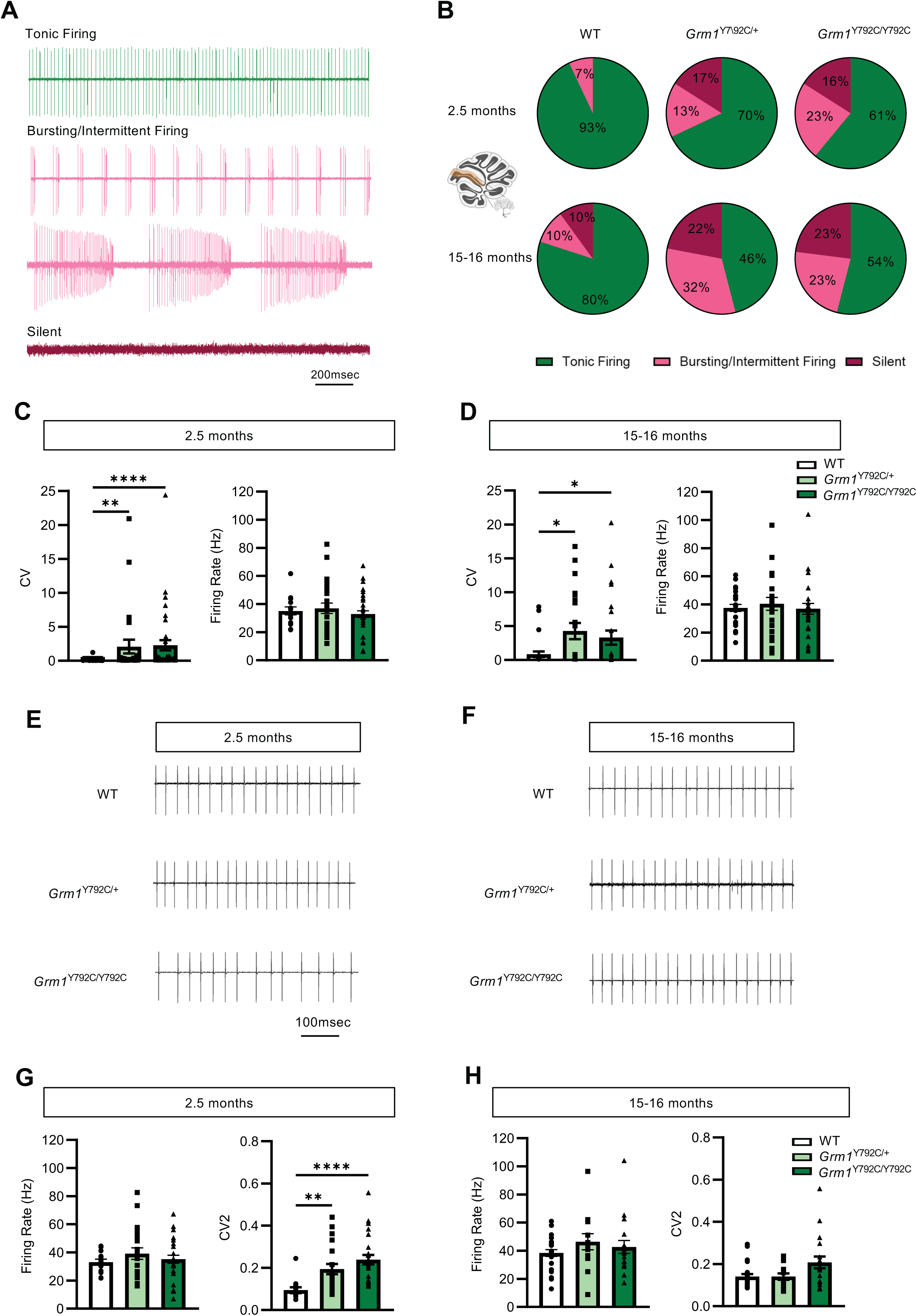
Spontaneous Purkinje cell firing pattern is disrupted in *Grm1* mutant mice. **(A)** Example traces from a tonic firing (green), bursting/intermittent firing (pink) and silent Purkinje cell (magenta). **(B)** The percentage of Purkinje cells displaying different modes of spontaneous firing activity recorded at 2.5 months (top) and 15-16 months of age (bottom) in lobule V. **(C)** Quantification of the coefficient of variation (CV) of the interspike interval, and the mean firing rate of all firing Purkinje cells at 2.5 months of age. CV: WT vs *Grm1*^Y792C/+^: *P*=0.0043, WT vs *Grm1*^Y792C/Y792C^: *P*<0.0001, *Grm1*^Y792C/+^ vs *Grm1*^Y792C/Y792C^: *P=*0.2895; mean firing rate: WT vs *Grm1*^Y792C/+^: *P*=0.9262, WT vs *Grm1*^Y792C/Y792C^: *P*=0.8846, *Grm1*^Y792C/+^ vs *Grm1*^Y792C/Y792^: *P*=0.5459. n=14 (WT), n=25 (*Grm1*^Y792C/+^), n=37 (*Grm1*^Y792C/Y792C^). **(D)** Quantification of the coefficient of variation (CV) of the interspike interval, and the mean firing rate of all firing Purkinje cells at 15 months of age. CV: WT vs *Grm1*^Y792C/+^: *P*=0.0433, WT vs *Grm1*^Y792C/Y792C^: *P*=0.0163, *Grm1*^Y792C/+^ vs *Grm1*^Y792C/Y792C^: *P*>0.9999; mean firing rate: WT vs *Grm1*^Y792C/+^: *P*>0.9999, WT vs *Grm1*^Y792C/Y792C^: *P*>0.9999, *Grm1*^Y792C/+^ vs *Grm1*^Y792C/Y792C^: *P*=0.8172. n=27 (WT), n=22 (*Grm1*^Y792C/+^), n=27 (*Grm1*^Y792C/Y792C^). **(E, F)** Example traces of tonic firing Purkinje cell recordings at 2.5 months (E) and 15-16 months (F). **(G)** Mean firing rate and precision measured using CV2 of the interspike interval of tonic firing Purkinje cell at 2.5 months of age. Mean firing rate: WT vs *Grm1*^Y792C/+^: *P*=0.4916, WT vs *Grm1*^Y792C/Y792C^: *P*=0.9120, *Grm1*^Y792C/+^ vs *Grm1*^Y792C/Y792C^: *P*=0.6348; CV2, WT vs *Grm1*^Y792C/+^: *P*=0.0036, WT vs *Grm1*^Y792C/Y792C^: *P*<0.0001, *Grm1*^Y792C/+^ vs *Grm1*^Y792C/Y792C^: *P*=0.4365. n=13 (WT), n=21 (*Grm1*^Y792C/+^), n=27 (*Grm1*^Y792C/Y792C^). **(H)** Mean firing rate and precision measured using CV2 of the interspike interval of tonic firing Purkinje cell at 15-16 months of age. Mean firing rate: WT vs *Grm1*^Y792C/+^: *P*=0.6423, WT vs *Grm1*^Y792C/Y792C^: *P*>0.9999, *Grm1*^Y792C/+^ vs *Grm1*^Y792C/Y792C^: *P*=0.7577; CV2: WT vs *Grm1*^Y792C/+^: *P*>0.9999, WT vs *Grm1*^Y792C/Y792C^: *P*=0.1459, *Grm1*^Y792C/+^ vs *Grm1*^Y792C/Y792C^ *P*=0.2882. n=24 (WT), n=13 (*Grm1*^Y792C/+^), n=19 (*Grm1*^Y792C/*Y792C*^). Statistical significance was determined by one-way ANOVA followed by Tukeys’ multiple comparison test, or Kruskal-Wallis test followed by Dunn’s multiple comparison test. Error bars represent SEM. \**P*<0.05, \*\**P*<0.01, \*\*\*\**P*<0.0001. For full statistical data, see Supplementary Table 2.

At the early stage of the disease, the majority (93%) of WT Purkinje cells displayed tonic firing, while the remaining 7% exhibited bursting/intermittent firing. In contrast, the proportion of tonically firing cells was reduced to 70% and 61% in *Grm1*^Y792C/+^ and *Grm1*^Y792C/Y792C^ mice, respectively (Fig. 6B). An increased proportion of the *Grm1* mutant Purkinje cells displayed bursting/intermittent firing activity, which was reflected by larger number of *Grm1* mutant Purkinje cells displaying higher CV values (Fig. 6B, C). The remaining *Grm1* mutant Purkinje cells remained silent with no firing activity (Fig. 6B, C). We found no differences in the mean firing rate of all firing cells (tonic plus intermittent firing/bursting cells) between WT and *Grm1* mutant mice (WT vs *Grm1*^Y792C/+^: *P*=0.9262, WT vs *Grm1*^Y792C/Y792C^: *P*=0.8846) (Fig. 6C).

We made similar observations at late stage of the disease, where the proportion of tonically firing Purkinje cells remained lower in *Grm1* mutant mice at 46% (*Grm1*^Y792C/+^) and 54% (*Grm1*^Y792C/Y792C^), compared to 80% of WT Purkinje cells (Fig. 6B). An increased number of mutant Purkinje cells displayed bursting/intermittent firing compared to WT, which was reflected by an increased number of mutant Purkinje cells displaying higher CV values (Fig. 6B, D). Moreover, we observed an increased percentage of silent Purkinje cells in mice from both *Grm1* mutant genotypes. Similar to the early disease stage, the mean firing rate of all firing Purkinje cells in both *Grm1* mutant mice was similar to WT mice (Fig. 6D).

We next investigated the tonic firing rate and regularity in *Grm1* mutant Purkinje cells. At the early disease stage, the mean firing rate of tonic firing Purkinje cells was similar in mutant and WT mice (Fig. 6E, G). However, Purkinje cells of both *Grm1* mutant mice fired more irregularly and with decreased precision as reflected by a larger CV2 value, which measures the regularity of adjacent interspike intervals (WT vs *Grm1*^Y792C/+^: *P*=0.0036, WT vs *Grm1*^Y792C/Y792C^: *P*<0.0001) (Fig. 6G). However, at the late stage of the disease, the mean firing rate and regularity of tonic firing PCs of *Grm1* mutant mice were similar to WT mice (Fig. 6E, H). These results demonstrate that the spontaneous activity of Purkinje cells is disrupted early in both heterozygous and homozygous *Grm1* mutants.

### Lobule-specific variations in disrupted spontaneous activity of PCs

Given the region-specific vulnerability of Purkinje cells in cerebellar ataxia^36,41,42^ and the fact that Purkinje cell activity differs between cerebellar modules^8,43^, we next set out to investigate whether enhanced mGluR1 signalling might impact on the spontaneous activity of Purkinje cells in a lobule-specific manner. We recorded spontaneous activity of Purkinje cells specifically in lobule III and lobule X at 2.5 months of age. As above (Fig. 6), we found that the percentage of tonically firing Purkinje cells was reduced in *Grm1* mutant mice. In lobule III, 78% of in *Grm1*^Y792C/+^ and 55% of the *Grm1*^Y792C/Y792C^ Purkinje cells exhibited tonic firing, compared to 92% in WT mice. 32% of *Grm1*^Y792C/Y792C^ Purkinje cells exhibited bursting activity, compared to 11% in *Grm1*^Y792C/+^ and 8% in WT mice and as indicated by higher CV values (Fig. 7A, B). The remaining mutant Purkinje cells were silent. Despite the differences in firing patterns, the mean firing rate across all firing cells (tonic and intermittent firing/bursting cells) did not differ significantly between WT and *Grm1* mutant mice (WT vs *Grm1*^Y792C/+^: *P*=0.4041, WT vs *Grm1*^Y792C/Y792C^*: P*=0.5726, *Grm1*^Y792C/+^ vs *Grm1*^Y792C/Y792C^*: P*=0.9417) (Fig. 7B).

**Figure 7.**
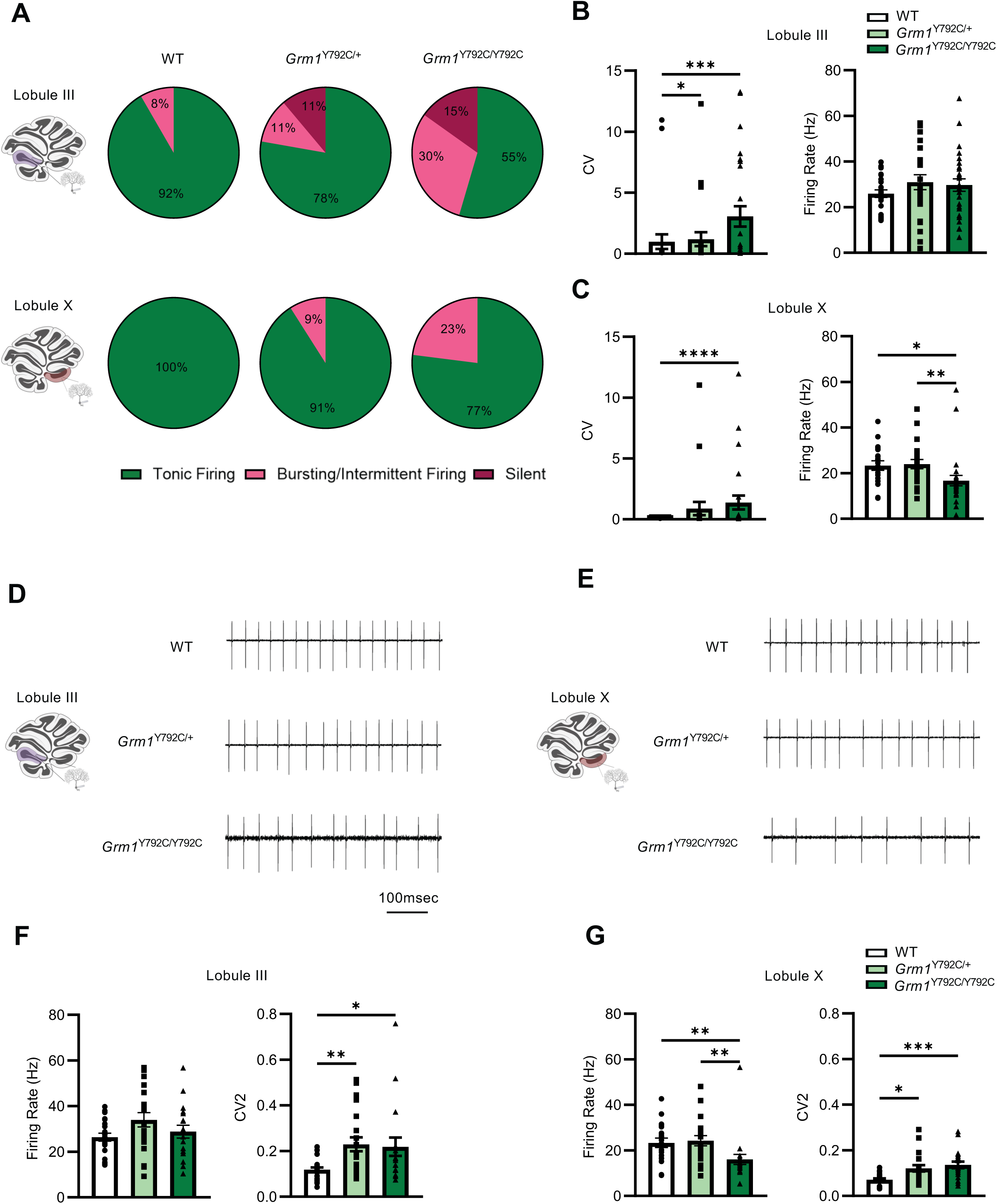
Lobule-specific disruptions in spontaneous firing patterns of Purkinje cells in *Grm1* mutant mice. **(A)** The percentage of Purkinje cells displaying different modes of spontaneous firing activity recorded from lobule III (top) and lobule X (bottom) from 2.5-month-old WT, *Grm1^Y792C/+^* and *Grm1^Y792C/Y792C^* mice. (**B)** The coefficient of variation (CV) of the interspike interval, and the mean firing rate of all firing Purkinje cells from lobule III. CV: WT vs *Grm1*^Y792C/+^: *P*=0.0343, WT vs *Grm1*^Y792C/Y792C^: *P*=0.0009, *Grm1*^Y792C/+^ vs *Grm1*^Y792C/Y792C^: *P*=0.9458; mean firing rate: WT vs *Grm1*^Y792C/+^: *P*=0.4041, WT vs *Grm1*^Y792C/Y792C^ *P*=0.5726, *Grm1*^Y792C/+^ vs *Grm1*^Y792C/Y792C^: *P*=0.9417. n=24 (WT), n=24 (*Grm1*^Y792C/+^), n=28 (*Grm1*^Y792C/Y792C^). **(C)** The coefficient of variation (CV) of the interspike interval, and the mean firing rate of all firing Purkinje cells from lobule X. CV: WT vs *Grm1*^Y792C/+^: *P*=0.0689, WT vs *Grm1*^Y792C/Y792C^: *P*<0.0001, *Grm1*^Y792C/+^ vs *Grm1*^Y792C/Y792C^: *P*=0.1224; mean firing rate: WT vs *Grm1*^Y792C/+^: *P*>0.9999, WT vs *Grm1*^Y792C/Y792C^: *P*=0.0105, *Grm1*^Y792C/+^ vs *Grm1*^Y792C/Y792C^: *P*=0.0050. n=19 (WT), n=22 (*Grm1*^Y792C/+^), n=26 (*Grm1*^Y792C/^Y792C). (**D, E)** Example traces of tonic firing Purkinje cell recordings from lobule III (D) and lobule X (E) of 2.5-month-old WT, *Grm1*^Y792C/+^ and *Grm1*^Y792C/Y792C^ mice. **(F)** Mean firing rate and precision measured using CV2 of the interspike interval of tonic firing Purkinje cells from lobule III. Mean firing rate: WT vs *Grm1*^Y792C/+^: *P*=0.0932, WT vs *Grm1*^Y792C/Y792C^: *P*=0.7884, *Grm1*^Y792C/+^ vs *Grm1*^Y792C/Y792C^: *P*=0.3637; CV2: WT vs *Grm1*^Y792C/+^: *P* =0.0066, WT vs *Grm1*^Y792C/Y792C^ :*P*=0.0348, *Grm1*^Y792C/+^ vs *Grm1*^Y792C/Y792C^: *P*>0.9999. n=22 (WT), n=21 (*Grm1*^Y792C/+^), n=18 (*Grm1*^Y792C/^Y792C). **(G)** Mean firing rate and precision measured using CV2 of the interspike interval of tonic firing Purkinje cells from lobule X. Mean firing rate: WT vs *Grm1*^Y792C/+^: n=20, *P*>0.9999, WT vs *Grm1*^Y792C/Y792C^: *P*=0.0088, *Grm1*^Y792C/+^ vs *Grm1*^Y792C/Y792C^: *P*=0.0051; CV2: WT vs *Grm1*^Y792C/+^: *P*=0.0137, WT vs *Grm1*^Y792C/Y792C^: *P*=0.0007, *Grm1*^Y792C/+^ vs *Grm1*^Y792C/Y792C^: *P*>0.9999. n=19 (WT), n=20 (*Grm1*^Y792C/+^), n=20 (*Grm1*^Y792C/Y792C^). Statistical significance was determined by one-way ANOVA followed by Tukeys’ multiple comparison test, or Kruskal-Wallis test followed by Dunn’s multiple comparison test. Error bars represent SEM. \**P*<0.05, \*\**P*<0.01, \*\*\**P*<0.001, \*\*\*\**P*<0.0001. For full statistical data, see Supplementary Table 2.

In lobule X, we also observed an increase in the proportion of bursting Purkinje cells in the *Grm1* mutants compared to WT Purkinje cells, which were all tonic firing (Fig. 7A, C). In contrast to lobule III, none of the *Grm1* mutant Purkinje cells were silent in lobule X. Furthermore, we found that the mean firing rate across all firing Purkinje cells (tonic and intermittent firing/bursting cells) was significantly lower in lobule X of the homozygous *Grm1*^Y792C/Y792C^ mice compared to both heterozygous *Grm1*^Y792C/+^ and WT mice (WT vs *Grm1*^Y792C/Y792C^: *P*=0.0105, *Grm1*^Y792C/+^ vs *Grm1*^Y792C/Y792C^: *P*=0.0050) (Fig. 7C).

We next examined the properties of tonic firing Purkinje cells in lobule III and lobule X. The mean firing rate of tonic firing Purkinje cells of lobule III was not significantly different between WT, *Grm1*^Y792C/+^ and *Grm1*^Y792C/Y792C^ mice (Fig. 7D, F). In contrast, in lobule X, the mean firing rate of *Grm1*^Y792C/Y792C^ Purkinje cells was significantly lower than in WT and *Grm1*^Y792C/+^ (WT vs *Grm1*^Y792C/+^: *P*>0.9999, WT vs *Grm1*^Y792C/Y792C^: *P*=0.0088, *Grm1*^Y792C/+^ vs *Grm1*^Y792C/Y792C^: *P*=0.0051) (Fig. 7D, G). We found that regardless of lobule location, tonic firing Purkinje cells of *Grm1* mutant mice fired more irregularly and with decreased precision, as reflected by the larger CV2 value (Fig. 7E, G).

Together, these findings demonstrate that gain-of-function mGluR1 signalling modulates Purkinje cell activity in a region-dependent manner that is also dependent on the level of mGluR1 overactivity.

## Discussion

The mGluR1 signalling cascade is critical to the normal functioning of Purkinje cells and its disruption results in cerebellar ataxia in humans and animal models. However, the pathophysiological role of aberrant mGluR1 signalling in SCA remains enigmatic as both increase and decrease of mGluR1 function have been linked to cerebellar disease^4,5^. Increasing evidence points to the presence of enhanced mGluR1 function during early stages of disease. For example, early-stage mouse models of SCA1 and SCA2 display overactive mGluR1 currents^17,18^. However, changes in other ion channels have also been reported in these models^44,45^, complicating the interpretation of results. As a consequence, it remains unclear whether increased mGluR1 activity represents a compensatory change or disease-driving mechanism in SCA. In this study, we sought to address the controversy around altered mGluR1 activity by using a mouse model that harbours a gain-of-function mutation in mGluR1, which we had previously identified in SCA44 patients^22^. We demonstrate that the p.Y792C mutation indeed behaves as a gain-of-function mutation in Purkinje cells, resulting in enhanced mGluR1 currents. We show that *Grm1* mutant mice exhibit cardinal features of SCA, including progressive motor coordination deficits, reduced climbing fibre territory, and Purkinje cell firing deficits^40,46^. These findings demonstrate for the first time that enhanced mGluR1 function in Purkinje cells is a direct driver of cerebellar dysfunction and disease.

We found that homozygous *Grm1*^Y792C/Y792C^ mutants displayed a more severe motor phenotype with progressive motor deficits manifesting from three months of age. Motor deficits in the heterozygous *Grm1*^Y792C/+^ animals were more subtle and only detectable from 12 months onwards on the narrow balance beam. Similarly, Purkinje cell soma atrophy was detected earlier in homozygous compared to heterozygous mutants. These findings suggest a gene-dosage effect of gain-of-function *Grm1* on phenotypic expression. Homozygous SCA44 patients have not been described, but gene dosage effects and resulting earlier age of onset and more severe clinical phenotypes have been reported in rare homozygous SCA3 and SCA6 patients^47,48^.

mGluR1 is widely expressed in the central nervous system, and loss of mGluR1 function results in behavioural phenotypes that are associated with broader neurological dysfunction including ataxia and impaired motor learning^49^ but also context-dependent deficits in associative learning^50^, and disrupted prepulse inhibition^33^. Similarly, *GRM1* loss-of-function mutations in humans cause SCAR13^10,11^, a rare neurodevelopmental disorder characterised by ataxia and intellectual disability, and have also been linked to schizophrenia^51^. In contrast, in our gain-of-function *Grm1* mutants, we only observed motor phenotypes consistent with cerebellar dysfunction and did not find deficits in any of the non-cerebellar behaviours assessed. These findings suggest that the cerebellum is particularly vulnerable to increased mGluR1 function, which may be due to the lack of compensatory mechanisms that are present in other brain regions. One such compensatory mechanisms might be the expression of other group I mGlu receptors; while mGluR1 and mGluR5 are co-expressed in many brain regions where they engage in complex interactions^52^, their expression in Purkinje cells is mutually exclusive^53^.

We identified deficits in spontaneous Purkinje cell activity in both heterozygous and homozygous *Grm1* mutants at an early age, highlighting the critical role of mGluR1 function in spontaneous Purkinje cell activity. Moreover, our findings are consistent with the hypothesis that Purkinje cell firing abnormalities are early drivers of cerebellar dysfunction. We observed a reduction in tonic-firing Purkinje cells and concomitant increase in bursting/intermittent firing and silent Purkinje cells in *Grm1* mutant mice. Silent Purkinje cells are likely depolarised and in a depolarisation block^54,55^. Our findings share important parallels with observations in other mouse mutants in which metabotropic glutamatergic signalling is enhanced. Increased activity of the mGluR1-coupled TRPC3 ion channel has been shown to increase the proportion of silent Purkinje cells^55^, while knockout of the EAAT4 glutamate transporter that maintains low synaptic glutamate levels in Purkinje cells results in an increased proportion of bursting Purkinje cells^56^. PKCy, which acts downstream of mGluR1, reduces large calcium-activated potassium channel currents^57^, which has been shown to contribute to silent Purkinje cells^54^ and firing irregularity in tonic-firing Purkinje cells^58^. It will be interesting to further dissect the downstream mechanisms by which enhanced mGluR1 disrupts Purkinje cell activity and to identify points for potential intervention.

Our study sheds light on the important question of regional vulnerability of Purkinje cell subpopulations in the cerebellum in disease. Patterned cell death has been observed in a number of cerebellar mouse mutants^59^, and prominent anterior vermis degeneration occurs in ataxia patients and mouse models of SCA1^36^ and ARSACS^37^. This region-specific vulnerability is thought to be underpinned by distinct molecular identities of Purkinje cell subpopulations. Purkinje cells express aldolase C, also known as zebrin, as well as other molecules in a striped pattern, with anterior lobules containing predominantly zebrin-negative Purkinje cells while posterior lobules are enriched in zebrin-positive Purkinje cells^60^. Furthermore, the regional identity of Purkinje cells is also associated with functional differences and correlates with differences in firing frequency, intrinsic plasticity, and synaptic transmission^8,43,61–63^. Notably, we found that the impact of enhanced mGluR1 activity also manifests in a lobule-dependent manner. Purkinje cell soma atrophy and reduced climbing fibre innervation were observed early in anterior lobules (III and V) and only later in the posterior lobule X. Moreover, we observed a larger decrease in tonic-firing Purkinje cells in lobules III and V compared to lobule X, with silent cells detected only in anterior lobules. These results support the interesting hypothesis that anterior Purkinje cells are more vulnerable to the pathogenic effects of increased mGluR1 activity. Several factors may contribute to this. mGluR1 and components of its signalling cascade are differentially expressed in different cerebellar lobules. For example, expression of the mGluR1 receptor itself as well as the downstream effector TRPC3 channel are greater in the anterior lobules^8,61^. Specifically, the short and more active TRPC3c isoform is enriched in anterior lobules^61^, likely resulting in larger intracellular calcium levels in zebrin-negative Purkinje cells. Glutamate clearance also differs across cerebellar lobules, with the glutamate transporter EAAT4 being highly expressed around zebrin-positive cells, promoting faster glutamate clearance and limiting mGluR1 activation^56,64^. This contrasts with the slower glutamate clearance around zebrin-negative Purkinje cells due to lower EAAT4 expression, resulting in prolonged mGluR1 activation and likely exacerbating the deleterious effects of hyperactive mGluR1 signalling. These factors could collectively enhance intrinsic excitability and susceptibility of anterior-lobule Purkinje cells, resulting in early degenerative changes. The Purkinje cell soma atrophy observed in the *Grm1* mutant mice might be an adaptive, compensatory mechanism in response to the altered excitability of the Purkinje cells^54^. The observed decrease in tonic Purkinje cell firing rate in lobule X at 3 months might also reflect an early adaptive compensatory change and could contribute to the greater resilience of this region. Additional compensatory changes in the *Grm1* mutant mice together with an age-related decline in firing precision in the WT animals might explain the similar levels of firing precision in tonic-firing WT and mutant Purkinje cells that we observed at late-disease stage. Although we identified disease-relevant cellular and behavioural phenotypes in the *Grm1* mutant animals that worsened with age, we did not observe any Purkinje loss. Moreover, the observed disease phenotypes in the heterozygous *Grm1*^Y792C/+^ animals were subtle and differ from the apparent ataxia and cerebellar atrophy reported in heterozygous SCA44 patients^22^. These observations are similar to findings in other mouse models of SCA, particularly those that express disease mutations at physiological levels and display much subtler phenotypes compared to highly overexpressing transgenic mouse models^27,65,66^. The discrepancy between human and mouse pathology may relate to the limited lifespan of mice, which may not be long enough to complete the full cascade of pathological changes that follows the initiating event^67^. Furthermore, human cerebellar Purkinje neurons exhibit non-allometric structural expansions compared to mice including increased dendritic spine density and parallel fibre input, as well as a non-allometric distribution of ion-channel density^68^. These species differences may render human Purkinje cells more vulnerable to excitotoxicity over a lifespan. The more subtle changes detected in the *Grm1* mutant animals, particularly in the heterozygous *Grm1*^Y792C/+^ mice, likely represent early changes during disease. Hence, these animal models may be particularly valuable for investigating early interventions that could prevent further progression of pathology.

Many of the phenotypes we report in the *Grm1* mutant mice including Purkinje cell firing deficits and climbing fibre abnormalities have been demonstrated in other mouse models of SCA including SCA1 and SCA2^39,40,46^. Together with the studies that report enhanced mGluR1 currents in these models^17,18^, these findings suggest that enhanced mGluR1 activity represents a common disease mechanism in genetically distinct SCAs. Thus, pharmacological inhibition of mGluR1 presents a promising therapeutic strategy at least for a subgroup of SCAs.

## Methods

### Animals and Husbandry

All procedures and experiments were carried out in accordance with the Animals (Scientific Procedures) Act 1986, UK, amendment Regulations 2012 (SI 4 2012/3039). All mice were group-housed and kept under a 12-hour day/night cycle with food and water available *ad libitum* in a controlled environment maintained at a temperature of 22 ± 2°C with a humidity level of 55 ± 10%.

### Generation of *Grm1* mutant mice and genotyping

*Grm1* mutant mice were generated under Genome Editing Mice for Medicine (GEMM) programme at the Mary Lyon Centre, Medical Research Council Harwell, UK. Mutant animals were generated by pronuclear injection into one-cell stage embryos (background: C57BL/6NTac) with Cas9 mRNA, single guide mRNA (protospacer sequence: GCAGGTGGTGTACATAGTGA, PAM sequence AGG) and single-stranded oligonucleotide (sequence CTACTATGCCTTCAAGACCCGCAACGTGCCGGCCAATTTCAATGAGGCTAAATACATCGCgTTCACTATGTgCACCACCTGCATCATCTGGCTGGCTTTTGTTCCCATTTACTTTGGGAGCAACTACAAGAT) in microinjection buffer (10 mM Tris–HCl, 0.1 mM EDTA, 100 mM NaCl, pH7.5). This introduced the SCA44 point mutation (A>C; p.Tyr792Cys) in the endogenous *Grm1* gene as well as an additional silent base pair substitution (C>G; p.Ala782Ala) in the nearby PAM site to prevent re-cutting by the Cas9 enzyme. The MGI ID for the generated allele (Grm1^em1H^) is MGI:6451428.

Routine genotyping was performed from ear tissue samples using PCR (Fwd: 5′-CTCCCATGCCCATTTTGTCC-3′; Rev: 5′-AAAAGCCAGCCAGATGATGC-3′). After amplification, PCR products were digested using FastDigest Bsp1407I restriction enzyme (Thermo Scientific, FD0933), which recognizes a TGTACA restriction site that is only present in the *Grm1* WT allele.

### Behavioural testing

Two cohorts of male mice were used for the longitudinal behavioural testing at three, six, 12 and 18 months of age (for animal numbers at each age, see Supplementary Table 1). Behavioural tests were performed in the same order, as follows: (1) open field; (2) light/dark box; (3) elevated zero maze; (4) forced alteration Y-maze, (5) Locotronic gait analysis, (6) MouseWalker gait analysis, (7) balance beam. The balance beam testing began only at 6 months of age. Fear conditioning and pre-pulse inhibition testing were carried out only at 6 months of age. A second cohort of naïve male mice was used for the balance beam testing at three months of age. Experimentally naïve female mice were used for the accelerating Rotarod testing, balance beam test, Locotronic, and MouseWalker gait analysis at three and 12 months of age. All behavioural experiments and related data analysis was done blinded. The order of animals for each behavioural test was randomised.

### MouseWalker gait analysis

Mice were placed and acclimatised to the dark behavioural room at least 30 minutes before the start of experiments. The room was dark, except for the lights of the arena and a computer screen. Each mouse was placed on a raised sealed transparent platform (10 cm *x* ∼1 m) lit up by green and red LED light. The walking surface of the arena reflects green LED light except from where the mouse paws touch the glass. The mouse was allowed to run in the apparatus for 1 min while its footsteps were filmed at 81 fps (Basler acA1300-75gc camera) and recorded using the StreamPix 7 software. Videos were analysed and gait parameters were extracted using a custom-trained DeepLabCut^69^ model and curated into csv files for analysis using custom-written Python scripts.

### Locotronic gait analysis

Paw misplacements during walking were assessed using a Locotronic apparatus (Intelli-bio, Seichamps, France). Mice were challenged to cross a horizontal ladder (bars 3 mm in diameter and 7 mm apart). The crossing was encouraged with a brighter start and a darker zone towards the end of the ladder. Mice were tested for three trials with 30-min interval between each trial. Trials where the mice took longer than 30s to complete the run were excluded from the analysis. The number of front and hind limb slips were analysed.

### Balance beam

The apparatus consisted of a beam (∼1m long) raised 50 cm above the ground and leading toa dark escape box. Each mouse performed three trials on a wide (25-mm diameter) and narrow (12.5-mm) beam, with a minimum of 10 min rest between trials. At the start of each trial, mice were positioned facing away from the escape box and allowed to turn around on the beam before traversing its length. The number of falls was recorded. Trials where the mouse did not attempt to turn or turned but stayed at the start of the beam for longer than 150s were excluded from the analysis. The percentage of mice traversing the beam without a fall was calculated.

### Rotarod

Mouse motor performance was evaluated using an accelerating Rotarod (Ugo Basile, Gemonio, Italy) that accelerated from 4 to 40 rpm over a 5-minute period. Mice were tested for three trials each day on four consecutive days and were allowed to rest for 15 min between trials. The trial ended when a mouse either fell from the rod or completed a passive double rotation. The latency to fall or complete passive double rotation was recorded for each trial for each mouse, and the mean value from each of the three trials was calculated to compute the latency to fall for each mouse each day.

### RNA extraction and RT-qPCR

Total RNA was extracted from three-month-old mouse cerebella using the RNeasy Mini kit (Qiagen) according to the manufacturer’s instructions and treated with DNase (Qiagen). 1µg of whole cerebellar RNA was synthesised into cDNA with the SuperScript III First-Strand Synthesis System (Invitrogen). qRT-PCR was performed using the LightCycler 480 SYBR Green master mix (Applied Biosystems, Thermo Scientific), and run on a StepOnePlus™ Real-Time PCR System (Applied Biosystems, Thermo Scientific) under the following reaction conditions: 95°C for 20 min, followed by 40 cycles at 95°C for 3s and 60°C for 30s. Primer sequences: *Grm1* (Fwd: 5’-CGCTCCAACACCTTCCTCAACATT-3’, Rev: 5’ GGGGTATTGTCCTCTTCCTCCACG-3’) and *Gapdh*(Fwd: 5’-TGTGTCCGTCGTGGATCTGA-3’, Rev: 5’-TTGCTGTTGAAGTCGCAGGAG-3’). All samples were analysed in triplicate, and *Grm1* expression levels were normalised against the expression levels of *Gapdh* (housekeeping gene) to give relative expression levels (2^-ΔCT^).

### Immunohistochemistry and imaging

Mice were anesthetized and transcardially perfused with 10% neutral buffered formalin (NBF). Cerebella were removed and postfixed with 10% NBF at 4°C, then cryoprotected with 30% sucrose in PBS at 4°C. Fixed tissues were embedded in OCT (Thermo Scientific) and cut on a cryostat into 30 µm-thick parasagittal sections mounted on SuperFrost Plus slides. Slides were washed in PBS for 10 min, incubated in 0.1M glycine for 30 min in a humidified chamber, incubated in blocking buffer (10% normal goat serum, 0.3% Triton-X 100 in PBS) for 1h at RT and incubated overnight at 4°C with the following primary antibodies in blocking buffer: rabbit anti-vesicular glutamate transporter 1 (vGluT1) (1:500; Synaptic Systems; 135303), rabbit anti-vesicular glutamate transporter 2 (vGluT2) (1:500, Synaptic Systems; 135403), guinea pig anti-calbindin D28K (1:500; Synaptic Systems; 214005), mGluR1a (1:1,000; Nittobo Medical; mGluR1a-GP-Af660). After incubation, slides were washed in blocking buffer three times, 10 min each, and incubated for 3 h at room temperature with the following secondary antibodies: AlexaFluor 488 goat anti-rabbit (1:500, Invitrogen, A32731) and AlexaFluor 594 goat anti-guinea pig (1:500, Invitrogen, A-11076). Slides were washed three times, 10 min each, in blocking buffer and once in PBS, before mounting in Vectashield mounting media (Vector Laboratories, H-1200). Images were acquired using an Axiovision fluorescent microscope (Zeiss) with 2.5x and 20x objective lenses. All quantifications were performed with 20x images using the “measure” function of Fiji/ImageJ software (version 2.9.0/1.53). At least 2-3 sections were quantified for each lobule (III, V and X) per mouse. Molecular layer thickness was measured as the distance from the top of the Purkinje cell soma to the top of the dendritic tree. CF territory was expressed as the height of the VGluT2-positive CF terminals (distance from the top of the Purkinje cell soma to the highest VGluT2 puncta in the molecular layer) divided by the molecular layer thickness. Density of Purkinje cells were measured by dividing the number of calbindin-positive Purkinje cells by the length of each section in the lobules. Calbindin-positive Purkinje cell somata were selected by “freehand selection” function of Fiji/ImageJ and their areas were measured.

### Western blotting

Frozen mouse cerebellar tissue was homogenised in RIPA buffer (Thermo Fisher Scientific) containing cOMPLETE protease inhibitor (Roche) and phosSTOP phosphatase inhibitor cocktail (Roche). Homogenates were incubated for 10 min at 4°C and centrifuged at 15,000 *g* for 30 min at 4°C to obtain protein supernatants. Protein concentration was determined using Pierce™ Bradford Protein Assay Kit (Thermo Fisher Scientific). 30 µg of protein lysate were resolved by SDS-PAGE using 4-12% NuPAGE™ Bis-Tris gel (Invitrogen) at 125V for 100 min and transferred to nitrocellulose membranes (Amersham) at 250 mA for 90 min. Membranes were blocked in 5% milk in TBS-T for 1 h at room temperature. Membranes were then incubated with the following primary antibody (diluted in TBS-T containing 3% BSA) overnight at 4°C: mouse anti-mGluR1α (1:2,500; BD Biosciences; 556389) and rabbit anti-β-actin (1:2,500; Abcam; ab179467). Membranes were washed in TBS-T three times, 10 min each, and incubated with anti-rabbit (1:5,000; Cytiva; NA934V) and anti-mouse (1:5000; Cytiva; NA931V) IgG horseradish peroxidase-conjugated secondary antibodies for 2 h at room temperature, followed by three 10-minute washes in TBS-T. Membranes were developed using Pierce™ ECL Western Blotting Substrate (Thermo Fisher Scientific) on a Bio-Rad ChemiDoc MP Imaging System. Densitometric analyses were performed using FIJI/ImageJ software (version 2.9.0/1.53). Band intensity was normalised to the intensity of beta-actin (loading control) on the same membrane to give relative expression levels.

### MRI methods

MRI imaging was performed as previously described^70^. *(Acquisitions)* Data were acquired on a 7T Biospec Bruker MRI scanner (Paravision 360.1), equipped with a mouse quadrature cryoprobe. Structural scans were acquired with a multi-gradient-echo sequence (75µm isotropic, TR=50 ms, TEs= [3 6.8 10.6] ms). A manganese chloride solution was used as a contrast agent. Therefore, mice were intraperitoneally injected with a 50 mg/kg solution of MnCl_2_ 24 hours prior to the scan. Twenty mice (10 *Grm1*^Y792C/Y792C^, 3 females, and 10 WT littermates, 1 female) were scanned at 2.5 and six months. Animals were induced with 4% isoflurane in oxygen and anaesthesia was maintained with 1-2% isoflurane in oxygen. Respiration was monitored with a breathing pillow. Temperature was monitored with a rectal probe. Isoflurane level was adapted to keep the animals at 80-110 rpm and a warm water-circulation heating pad was used to keep the animals at 36-37°C during scanning.

*(Analysis)* For each structural scan, individual echoes were denoised with nonlocal means^71^ and averaged together. Individual scans were then co-registered, applying first linear and affine transformations and then a series of nonlinear registrations. Information regarding global deformations (i.e. the overall brain sizes) are stored in these transformation models. The non-linear registration creates a vector field that maps every point in one image to another and provides information about localized deformation (i.e. the relative regional changes). In practice, data were analysed using the MBM Pydpider toolkit^72^ with a MAGeT registration for segmentation (MRI atlas)^70,73–77^. Group-wise registration was performed on the 40 images (20 animals, 2 time points). Regional volumes were extracted from the individual LSQ6 resampled atlases with the RMINC tools (R package).

### Cerebellar slice preparation for *ex vivo* electrophysiology

Older mice (15-16 months) were anaesthetized with an intraperitoneal injection of Phenobarbitone overdose, after which rapid transcardial perfusion with ice-cold NMDG aCSF cutting solution containing the following (in mM): 92 NMDG, 2.5 KCl, 1.25 NaH_2_PO_4_, 30 NaHCO_3_, 20 HEPES, 25 glucose, 2 thiourea, 5 Na-ascorbate, 3 Na-pyruvate, 0.5 CaCl_2_ and 10 MgSO_4_·7H_2_O with pH titrated to 7.3-7.4 with concentrated HCl (all Sigma-Aldrich) was performed before decapitation. Other mice were anaesthetized with 4% isoflurane and swiftly decapitated. For cell-attached recording experiments, extracted brains were chilled and parasagittal (250µm) cerebellar vermal slices were cut in ice-cold NMDG aCSF cutting solution. Slices were recovered in a hot NMDG aCSF solution (35°C) cutting solution for 10-12 min. After the initial recovery period, slices were transferred and held in HEPES holding aCSF at room temperature (RT) until recording, containing the following (in mM): 2 NaCl, 2.5 KCl, 1.25 NaH_2_PO_4_, 30 NaHCO_3_, 20 HEPES, 25 glucose, 2 thiourea, 5 Na-ascorbate, 3 Na-pyruvate, 2 CaCl_2_ and 2 MgSO_4_·7H_2_O with pH titrated to 7.3-7.4 with concentrated NaOH (all Sigma-Aldrich). For whole-cell mGluR1 recordings, brains were chilled and parasagittal (250µm) cerebellar vermal slices were cut with ice-cold sucrose solution cutting solution containing the following (in mM): 75 sucrose, 87 NaCl, 2.5 KCl, 1 NaH_2_PO4, 25 NaHCO_3_, 25 glucose, 6 MgCl_2_, 0.5 CaCl_2_. Slices were recovered in hot normal aCSF (35°C) containing the following (in mM) for 30 min: 126 NaCl, 2.5 KCl, 1 NaH_2_PO4, 26 NaHCO_3_, 2.4 CaCl_2_, 1.3 MgCl_2_, and 10 glucose (all Sigma-Aldrich). Slices were cooled to RT in aCSF before recording, containing the following (in mM) 126 NaCl, 3 KCl, 1 NaH_2_Po4, 26 NaHCO_3_, 2.4 CaCl_2_, 1.3 MgCl_2_ and 10 glucose (all Sigma-Aldrich).

### Extracellular cell-attached recordings

Extracellular cell-attached recordings were performed at 30-31°C in aCSF constantly bubbled with carbogen. Purkinje cells were visualised with DIC optics, lobules (III, V and X) were identified using 10x objectives, and individual Purkinje cells were identified with a water-immersion 40x objective. Recording glass patch pipettes (World Precision instruments, USA), of 4-6 MΩ resistance (PC-100, Narshinge, Japan), when filled with aCSF were used. Purkinje cell spontaneous activity was recorded for 10 min, with recordings between 6-9 min being used for analysis. The interspike interval (ISI) was measured and the mean firing rate (inverse of the mean ISI), coefficient of variation CV, and the local coefficient of variation CV2 were quantified using the following formulas, CV=^σ^(ISI)/^μ^(ISI) and CV2=2*|ISI_n_ - ISI_n-1_|/ (ISI_n+1_+ISI_n_).

### Whole cell electrophysiology

For whole-cell voltage-clamp recordings, recording pipettes were filled with internal solution containing (in mM): 122 K-methane sulfonate, 9 NaCl, 9 HEPES, 0.036 CaCl_2_, 1.62 MgCl_2,_ 0.18 EGTA, 4 Mg-ATP, 0.3 Tris-GTP, 14 Tris-creatine (all Sigma-Aldrich)^78^. Cells were held at -70mV, and series resistance compensation of up to 70% was made when possible. We excluded recordings with a series resistance greater than 30 MΩ. PF stimulation was adjusted to evoke a AMPA-mediated fast EPSCs with an amplitude of 450-600pA in the presence of GABA(A) receptor antagonist Gabazine (15 µM; Hellobio; HB0901). PF-mediated mGluR1 slow EPSC were evoked by stimulation of PFs (200Hz; 200-μs duration; 2-sweep average) with 2, 5, 10 or 20 pulses, and measured in the presence of AMPA/kainate receptor blocker NBQX (20 µM; Hellobio; HB0442) and glutamate transporter blocker TBOA (100 µM; Hellobio; HB0258). All electrophysiology data was acquired using a Multiclamp 700B amplifier (Molecular Devices, LLC., San Jose, CA), filtered at 4 kHz online, digitised at 20 kHz with ICT18 (Heka Instrument, Inc., Holliston, MA). Data acquisition and analysis were carried out using custom scripts in Igor pro software (WaveMetrics Inc., Portland, OR).

### Statistical analysis

For behavioural, electrophysiological, immunohistochemical and WB data, data were tested for normality using the Shapiro-Wilk normality test. We used parametric (t-test or one-/two-way ANOVA) for normally distributed data and nonparametric (Kruskal-Wallis or Mann-Whitney) statistical tests for data with non-normal distribution, followed by pairwise multiple comparison tests as indicated in the figure legends. Data are reported as mean±SEM. All figures and statistical analysis were computed using Prism v10 (GraphPad Software Inc., CA, USA). All statistical data are available in the Supplementary Information.

For MRI statistical analysis, cerebellar regional volumes were isolated using a hierarchical anatomical tree (RMINC package). A linear mixed effect model was used to assess the effect of age and genotype (fixed effects: age, genotype and sex; random intercept: mouse ID) on cerebellar regional volumes. Multiple comparisons were corrected with a false discovery rate (FDR) threshold of 10%. The interaction between age and genotype was not significant.

## Data Availability

The MRI data that support the findings of this study are available in Zenodo with DOI 10.5281/zenodo.14884600. Code is available on GitHub (https://github.com/clemoune/sca44-memri). All other data are available within the paper and its supplementary files.

## Acknowledgements

We thank members of the MLC Ward 6 and BMS Level 2 for maintenance of animal colonies. We would like to thank Greg Daubney, Manager at the Histology Small Research Facility, for assistance in the sectioning and immunostaining of paraffin-embedded blocks.

## Author contributions

Conceptualization: P.L.O., E.B.E.B., Methodology: M.F.I., E.M., P.L.O., E.B.E.B., Formal analysis: M.F.I, S.B., Y.C.C., C.L., R.S.B., Investigation: M.F.I, S.B., Y.C.C., C.L., Resources: J.P.L., E.M., P.L.O., E.B.E.B., Writing – Original Draft: M.F.I., E.B.E.B., Writing – Review & Editing: M.F.I, S.B., Y.C.C., C.L., R.S.B. J.P.L., E.M., P.L.O., E.B.E.B., Funding acquisition: J.P.L., P.L.O., E.B.E.B.

## Supplementary material

Supplementary material is available.

