## Supplemental Data for "Enhanced mGluR1 function causes motor deficits and region-specific Purkinje cell dysfunction"

### Supplementary Methods

#### Open field test

Open field test was used to assess general locomotor activity and anxiety-like behaviour in a novel environment[1]. Mice were placed in square arenas (40x40 cm) and allowed to freely explore the arena for 20 min. Light levels in the centre of the arena were 150-200 lux, between two and four mice were tested at a time, with one mouse per arena. EthovisionXT tracking software (Noldus, Netherlands) was used to track movements, and to calculate the total distance travelled (in 5-min time bins), and the time spent in the centre of the arena.

#### Light/dark box test

The light/dark box test was used to assess anxiety-like behaviour. Mice were placed in the light side of the box in the light-dark test apparatus (41(L) x 20(W) x 26(H) cm, length of light section: 25 cm) and allowed to freely explore the apparatus for 5 min. Mouse activity was tracked using Ethovision XT software (Noldus, Netherlands). The time spent and the frequency of entry in the light zone, the dark zone, and the entry zone (*i.e.*, the transition zone between the light and dark zones) were analysed.

#### Elevated zero maze

The elevated zero maze was used to assess anxiety-like behaviour in the mice. The maze inlcuded a circular arena, which was made up of two open sections and two closed sections, with each section about 30cm long. The mice were placed in an open section and allowed to explore the arena for 5 min. Their activity was tracked using Ethovision XT (Noldus, Netherlands). The time and frequency spent in the open and the closed sections were analysed.

#### Forced alteration Y-maze

Forced alteration Y-maze was used to test the spatial working memory of the mice. The Y-maze apparatus consisted of three clear Perspex arms (30 x 8 x 20 cm) arranged at 120° angles, and connected to a central zone The test was divided into two phases: the habituation phase and the test phase. In the habituation phase the mouse was placed in the start arm and allowed to enter the familiar arm, while the entry to the unfamiliar novel arm was blocked. Once the mouse exited the start arm and entered the familiar arm, its activity was recorded for 5 min. Afterwards, the mouse was returned to its home cage for 1 min, while the maze was cleaned. During the test phase, the mouse was returned to the start arm and allowed to explore both the familiar and novel arms for 2 min. The position of the familiar and the novel arm were randomised and balanced between mice to reduce room position (spatial) bias. Animals were tracked using Ethovision XT software (Noldus, Netherlands), and the frequency of arm entries and the total time spent in each arm was analysed. To assess the memory performance of the mice in this test, we calculated novel preference ratio from the test phase of the experiment using the formula: $Novel\ preference\ ratio\ = \frac{Time\ in\ novel\ arm\ (s)}{Time\ in\ novel\ arm\ (s) + Time\ in\ familiar\ arm\ (s)}$

**Fear conditioning**

This test was used to measure aversive learning and memory in the mice. A neutral conditioned stimulus (CS) – a tone, was paired with an aversive unconditioned stimulus (US) – a mild foot shock. After conditioning, we assessed the spatial context or the CS elicited fear in the absence of US which is shown as lack of movement (freezing). On day 1 – conditioning trial, each mouse was placed in Perspex boxes with metal grid floor and conditioned using a training protocol, consisting of a baseline period, audible tone (20 s), a single foot shock (0.4 mA for 1 s) that co-terminates with the tone, and a period with no stimuli at the end. On day 2 – context trial, each mouse was exposed to the same Perspex box with metal grid floor, but it was not exposed to shocks or tones. After 4 h, the mouse was exposed to a round arena with walls covered in white and black stripes and with Vanillin extract rubbed on the top. This way the mouse was exposed to a visually and olfactory new environment. In this environment, the tone (CS) was played to the mouse. Immobility (freezing) was used as a measure of learning/memory performance.

**Prepulse inhibition**

Prepulse inhibition of the mice was tested in soundproof boxes. The mice were exposed to initial set of 5 runs with no stimulus, just white noise, and then second set of 5 runs with the startle stimulus only (120 dB). This was followed by 10 of the three types of trials in a pseudorandom order: null stimulus; startle stimulus (0 dB followed by 56 dB; 0 dB followed by 58 dB; 0 dB followed by 68 dB); prepulse and startle stimulus (0 dB followed by 120 dB; 56 dB followed by 120 dB; 58 dB followed by 120 dB; 65 dB followed by 120 dB). The whole run was 45-50 min long. The percentage of prepulse inhibition (PPI%) was calculated for the startle stimulus at each prespulse tone using the formula:

$$PPI\% = 100 - (\frac{Prepulse\ tone}{Startle\ stimulus} x\ 100)$$

**Immunohistochemistry**

For 21-month-old mice (Supplementary Fig. 3), paraffin-embedded blocks were used. Coronal cerebellar sections were cut on a microtome at 10 μm and mounted on SuperFrost plus slides. Slides were de-paraffinized with two 5-minute washes in xylene and rehydrated by sequential 2-minute washes with 100%, 95%, 80%, and 70% ethanol. After another two 5-minute washes in PBS, slides were microwaved with antigen retrieval buffer (sodium citrate pH 6.0) for 20 min and allowed to cool for 30 min at RT. Slides were washed with PBS three times, 5 min each, and incubated with blocking buffer (PBS, 5% bovine serum albumin, 0.5% Tween-20) in a humidified chamber for 1 h at RT. After blocking, slides were incubated overnight at 4°C with guinea pig anti-calbindin D28K (1:500; Synaptic Systems; 214005). Slides were washed in blocking buffer three times, 5 min each, and incubated with AlexaFluor 594 goat anti-guinea pig secondary antibody (1:500, Invitrogen, A-11076) for 3 h at room temperature. Slides were washed in blocking buffer, 5 min each, and then once in PBS for 5 min, before mounting with Vectashield mounting media (Vector Laboratories, H-1200).

Immunostaining of 19-month-old mice (Supplementary Fig. 4) was performed as described in the main manuscript. Cerebellar sections were incubated with rabbit anti-vesicular glutamate transporter 1 (vGluT1) (1:500; Synaptic Systems; 135303) and guinea pig anti-calbindin D28K

(1:500; Synaptic Systems; 214005), followed by secondary antibodies AlexaFluor 488 goat anti-rabbit (1:500, Invitrogen, A32731) and AlexaFluor 594 goat anti-guinea pig (1:500, Invitrogen, A-11076).

**Supplementary Table 1.** Sample size for genotypes used for behavioural analyses at each experimental age. No animals were excluded from the analysis. No criteria were set for inclusion.

| | 3 Months | | 6 Months | 12 Months | 18 Months |
|---|---|---|---|---|---|
| **Male** | Cohort 1 | Cohort 2 | Cohort 1 | Cohort 1 | Cohort 1 |
| Wildtype | 15 | 10 | 14-15 | 14 | 5-8 |
| *Grm1*$^{Y792C/+}$ | 23 | 10 | 23 | 23 | 14-15 |
| *Grm1*$^{Y792C/Y792C}$ | 14 | 10 | 14 | 14 | 11-12 |
| **Female** | | | | | |
| Wildtype | 10 | | | 11 | |
| *Grm1*$^{Y792C/+}$ | 10 | | | 11-12 | |
| *Grm1*$^{Y792C/Y792C}$ | 10 | | | 11 | |

### Supplementary Figures

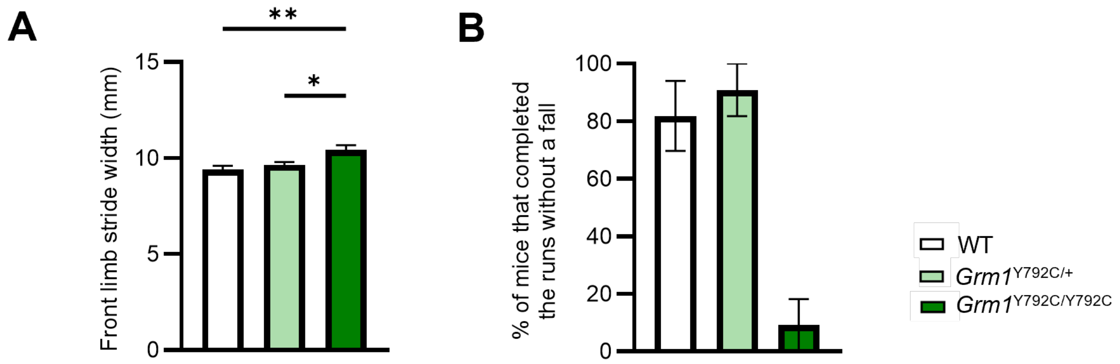


**Supplementary Figure 1. Gait and balance are disrupted in naïve 12-month-old female mutant *Grm1*$^{\mathbf{Y792C/Y792C}}$ mice. (A)** Quantification of the front limb stride width, *i.e.*, the distance between the right and the left front limb placement, assessed using the MouseWalker system. WT vs *Grm1*$^{Y792C/+}$: *P*=0.6905, WT vs *Grm1*$^{Y792C/Y792C}$: *P*=0.0029, *Grm1*$^{Y792C/+}$ vs *Grm1*$^{Y792C/Y792C}$: *P*=0.0231. Statistical significance was determined by one-way ANOVA followed by Tukey's multiple comparison test. For n-numbers, see Supplementary Table 1. **(B)** Quantification of the percentage of mice completing the runs on a wide (25 mm) balance beam without a fall. Error bars represent SEM. *$P<0.05$, **$P<0.01$, For full statistical data, see Supplementary Table 2.

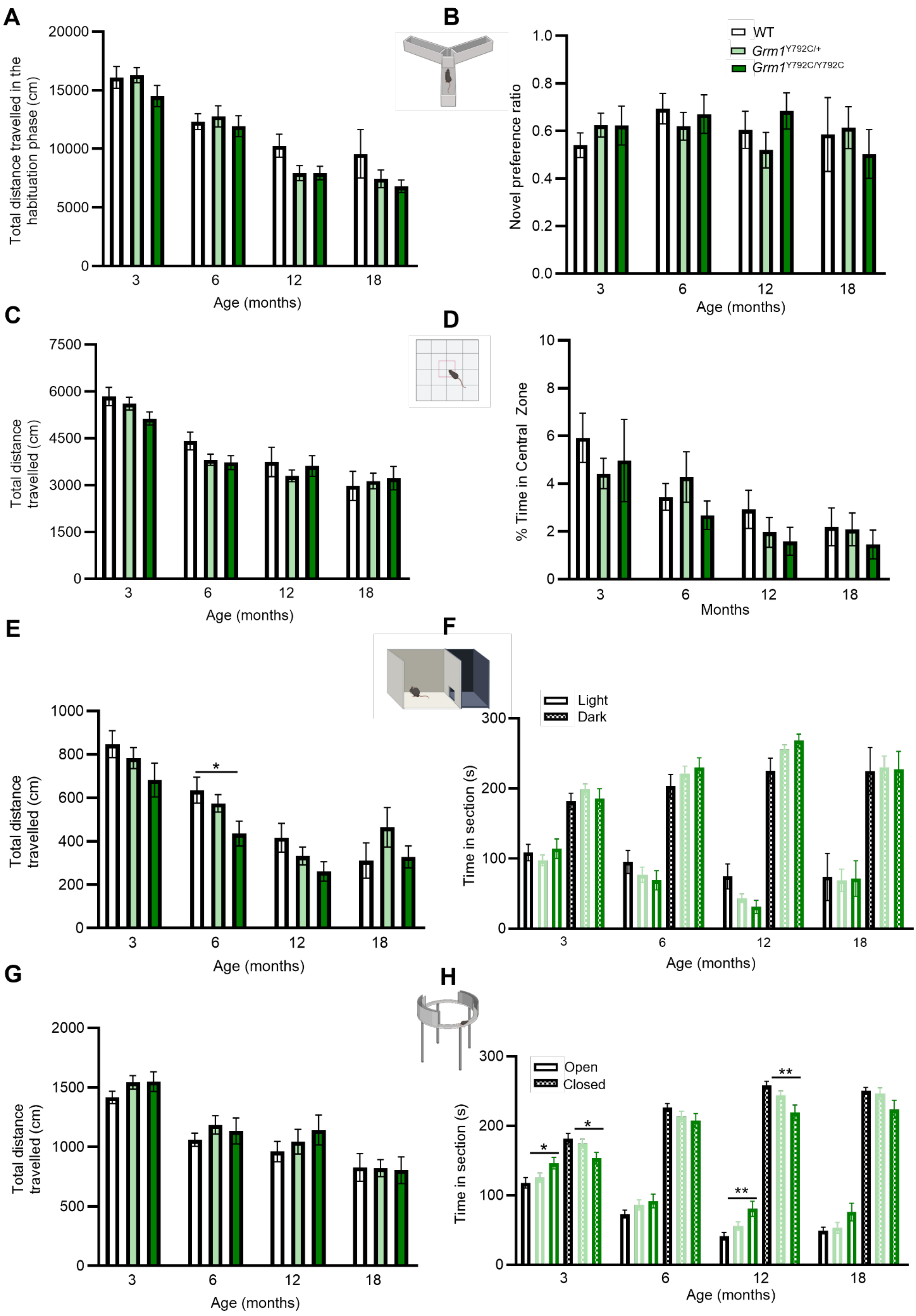
A
B
C
D
E
F
G
H
WT
Grm1Y792C/+
Grm1Y792C/Y792C
Total distance travelled in the habituation phase (cm)
Novel preference ratio
Total distance travelled (cm)
% Time in Central Zone
Time in section (s)
Age (months)
Months
Light
Dark
Open
Closed
3
6
12
18
*
**

**Supplementary Figure 2.** ***Grm1*** **mutant mice exhibit no deficits in non-motor behaviours**. **(A)** Total distance travelled by WT and *Grm1* mutant mice in the habitutation phase in the arms on Y-maze. **(B)** Time spent by WT and *Grm1* mutant mice in the novel arm on Y-maze compared to the total time spent in the novel and familiar arm expressed as the novel preference arm ratio. **(C)** Total distance travelled by WT and *Grm1* mutant mice in the open field. **(D)** Time spent in the central zone in open field assay by WT and *Grm1* mutant mice. **(E)** Total distance travelled by WT and *Grm1* mutant mice in the light/dark box. 6 months: WT vs *Grm1*$^{Y792C/+}$: *P*=0.6649, WT vs *Grm1*$^{Y792C/Y792C}$: *P*=0.037, *Grm1*$^{Y792C/+}$ vs *Grm1*$^{Y792C/Y792C}$: *P*=0.1376. **(F)** Time spent in the light (clear bars) versus dark (filled bars) areas in the light-dark box test by WT and *Grm1* mutant mice. **(G)** Total distance travelled by WT and *Grm1* mutant mice in the zero maze. **(H)** Time spent in the open and closed arms of the zero maze by WT and *Grm1* mutant mice. 3 months: time spent in the open arm, WT vs *Grm1*$^{Y792C/+}$: *P*=0.7499, WT vs *Grm1*$^{Y792C/Y792C}$: *P*=0.0436, *Grm1*$^{Y792C/+}$ vs *Grm1*$^{Y792C/Y792C}$: *P*=0.1221.Time spent in the closed arm,WT vs *Grm1*$^{Y792C/+}$: *P*=0.7499, WT vs *Grm1*$^{Y792C/Y792C}$: *P*=0.0436, *Grm1*$^{Y792C/+}$ vs *Grm1*$^{Y792C/Y792C}$: *P*=0.1221. 12 months: time spent in the open arm,WT vs *Grm1*$^{Y792C/+}$: *P*=0.3293, WT vs *Grm1*$^{Y792C/Y792C}$: *P*=0.0095, *Grm1*$^{Y792C/+}$ vs *Grm1*$^{Y792C/Y792C}$: *P*=0.3104. Time spent in the closed arm,WT vs *Grm1*$^{Y792C/+}$: *P*=0.3158, WT vs *Grm1*$^{Y792C/Y792C}$: *P*=0.0091, *Grm1*$^{Y792C/+}$ vs *Grm1*$^{Y792C/Y792C}$: *P*=0.3157. Statistical significance was determined by one-way ANOVA followed by Tukey's *post hoc* test or Kruskal-Wallis test followed by Dunn's multiple comparison test. Error bars represent SEM. *P*>0.05 when not indicated. **P*<0.05, ***P*<0.01. For full statistical data, see Supplementary Table 2.

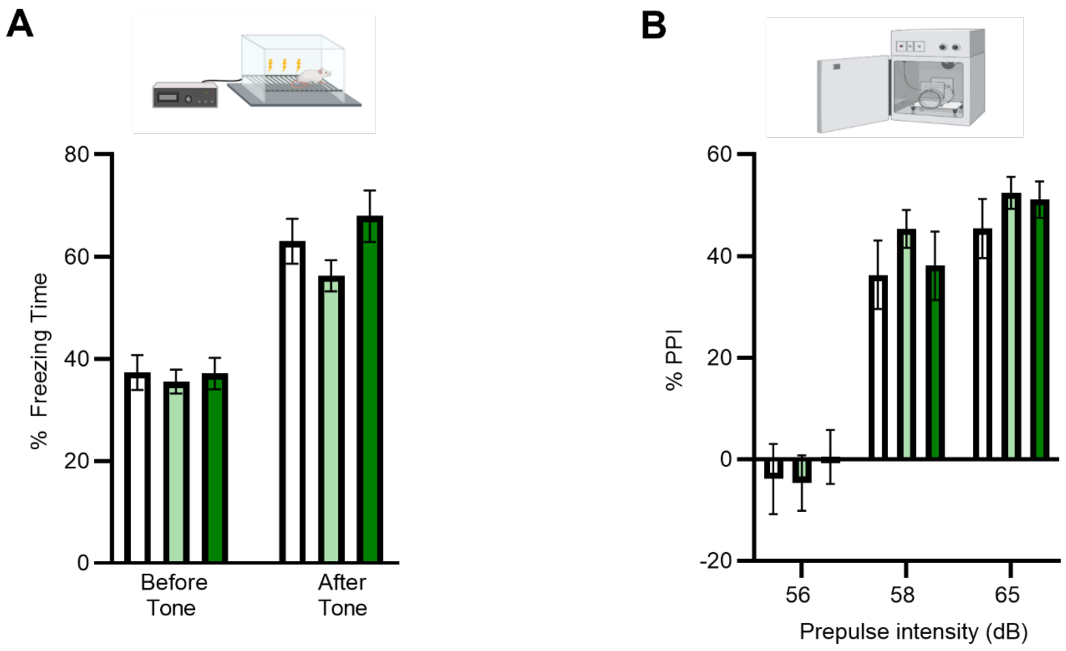


**Supplementary Figure 3. (A)** Percent freezing during context fear conditioning learning in six-month-old WT and *Grm1* mutant mice. (**B)** Prepulse inhibtion (PPI) values in six-month-old WT and *Grm1* mutant mice at three different prepule intensities. Statistical significance was determined by one-way ANOVA followed by Tukey's *post hoc* test or Kruskal-Wallis test followed by Dunn's multiple comparison test. For n-numbers, see Supplementary Table1. Error bars represent SEM. $P$>0.05 when not indicated. For full statistical data, see Supplementary Table 2.

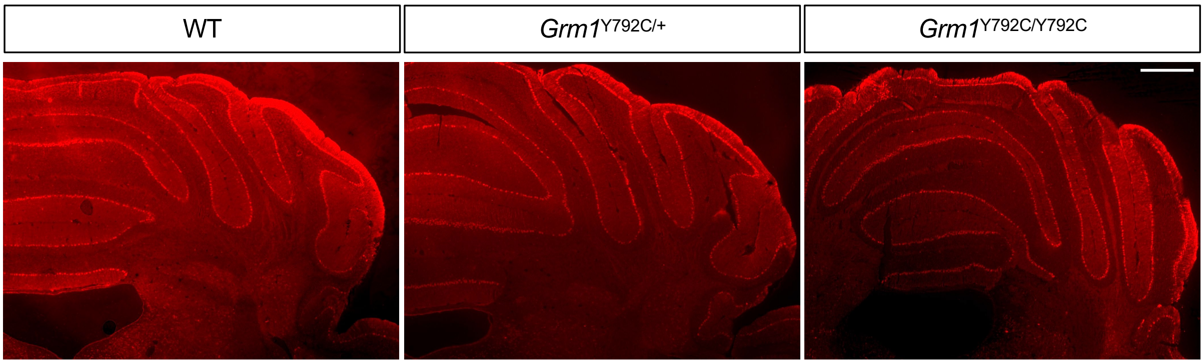


**Supplementary Figure 4. No cell loss was observed in 21-month-old WT and *Grm1* mutant mice.** Representative immunohistochemical staining for Purkinje cell marker Calbindin using coronal sections of 21-month-old WT and *Grm1* mutant cerebellum. Calbindin staining of mutant cerebellum was indistinguishable from WT. Scale bar: 500 µm.

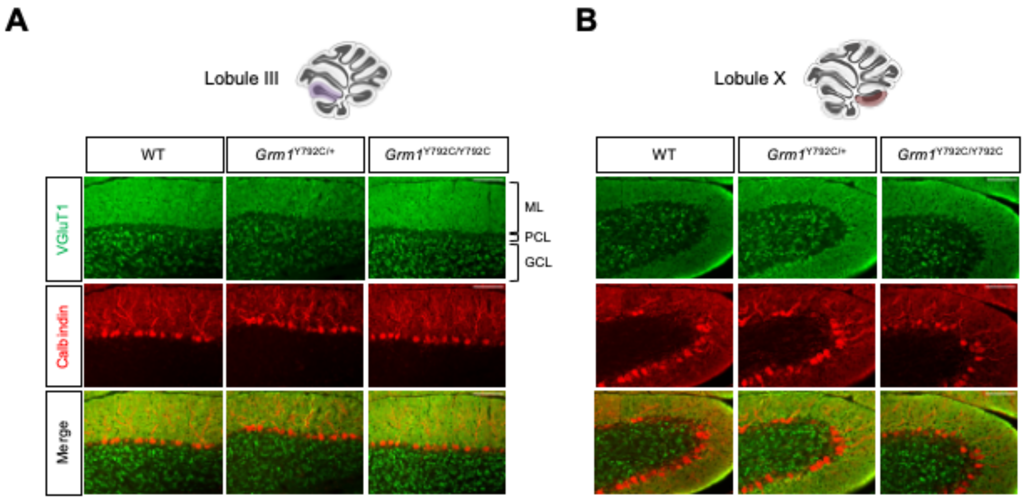


**Supplementary Figure 5. No changes in VGluT1 staining in *Grm1* mutant cerebellum.** Representative immunostaining for the parallel fibre synapse marker VGluT1 (green) and the Purkinje cell marker calbindin (red) in lobule III **(A)** and X **(B)** from 15-month-old WT and *Grm1* mutant cerebellum. ML: molecular layer; PCL: Purkinje cell layer; GCL: granule cell layer. Scale bar: 100 µm.

mGluR1

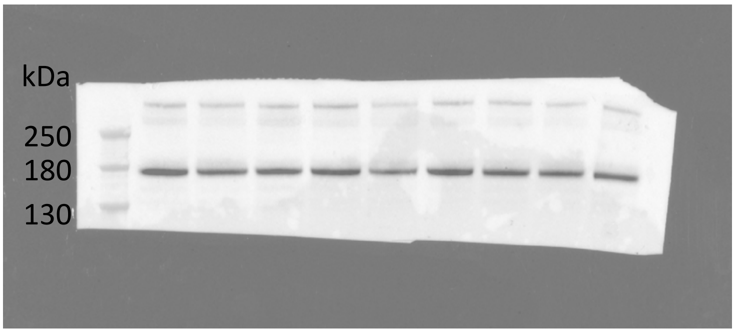


β-actin

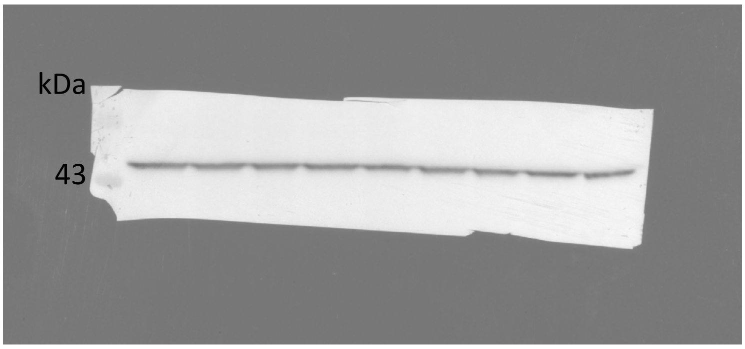


**Supplementary Figure 6. Original, uncropped images of Western blots shown in Figure 1E in the main manuscript.** Images are captured using the ChemiDoc Imaging System (Bio-rad). The same membrane is cut into different parts for incubation with primary antibodies against mGluR1 (upper blot) and β-actin as loading control (lower blot). The uncropped images are obtained by merging the colorimetric (for protein ladder) and chemiluminescence (for target protein band) images. The Color Prestained Protein Standard, Broad Range (NEB, P7719S) is used as molecular size marker.
